# Tunable metastability of condensates reconciles their dual roles in amyloid fibril formation

**DOI:** 10.1101/2024.02.28.582569

**Authors:** Tapojyoti Das, Fatima K. Zaidi, Mina Farag, Kiersten M. Ruff, Tharun Selvam Mahendran, Anurag Singh, Xinrui Gui, James Messing, J. Paul Taylor, Priya R. Banerjee, Rohit V. Pappu, Tanja Mittag

## Abstract

Stress granules form via co-condensation of RNA-binding proteins containing prion-like low complexity domains (PLCDs) with RNA molecules. Homotypic interactions among PLCDs can drive amyloid fibril formation that is enhanced by ALS-associated mutations. We report that condensation-versus fibril-driving homotypic interactions are separable for A1-LCD, the PLCD of hnRNPA1. Separable interactions lead to thermodynamically metastable condensates and globally stable fibrils. Interiors of condensates suppress fibril formation whereas interfaces have the opposite effect. ALS-associated mutations enhance the stability of fibrils and weaken condensate metastability, thus enhancing the rate of fibril formation. We designed mutations to enhance A1-LCD condensate metastability and discovered that stress granule disassembly in cells can be restored even when the designed variants carry ALS-causing mutations. Therefore, fibril formation can be suppressed by condensate interiors that function as sinks. Condensate sink potentials are influenced by their metastability, which is tunable through separable interactions even among minority components of stress granules.

## INTRODUCTION

Cytoplasmic stress granules (SGs) are inhomogeneous biomolecular condensates that form, in response to polysomal runoff caused by cellular stresses, via condensation. This combines reversible binding, oligomerization, phase separation, and percolation via a network of homotypic and heterotypic protein-protein, protein-RNA, and RNA-RNA interactions ^1–4^. SG formation is driven by a core protein-RNA interaction network involving mRNA, the RNA-binding proteins (RBPs) G3BP1/2, TIA1, CAPRIN1 and other RBPs including FUS, TDP-43, hnRNPA1, and hnRNPA2 ^1–3,5^. The pathogenesis of neurodegenerative and neuromuscular disorders such as amyotrophic lateral sclerosis (ALS), frontotemporal dementia (FTD), inclusion body myopathy (IBM), and multisystem proteinopathy (MSP) is associated with aberrant SG assembly and arrested disassembly ^6^. Cytoplasmic inclusions comprising insoluble deposits of SG proteins are among the defining pathological hallmarks of ALS, FTD and IBM ^7–12^.

Many mutations that lead to disease occur within intrinsically disordered, prion-like low-complexity domains (PLCDs) of RBPs. These mutations promote the formation of amyloid fibrils through homotypic interactions ^13–16^. This has led to the proposal that the interiors of SGs serve as crucibles of amyloid formation: the high local concentration of PLCDs within condensate interiors enables the crossing of threshold concentrations for fibril formation enabling homotypic, fibril-driving interactions ^17,18^. It has also been suggested that SGs are protective against fibril formation, because localization of TDP-43 to condensates other than SGs might be a key step in pathogenesis ^19^. The effects of condensate interfaces also add a novel nuance because interfaces accelerate fibril formation ^20–22^ via heterogeneous nucleation ^23^.

Taken together, one can envisage two distinct scenarios, one that integrates the effects of interfaces with the crucible hypothesis and an alternative that combines the effects of interfaces with the potentially protective effects of condensate interiors. Which of these two pictures applies to condensates is unclear. To adjudicate between the two scenarios, we investigated the underpinnings of condensate versus fibril formation in a system defined by homotypic interactions. Condensates and fibrils are disordered and ordered phases formed by the same proteins ^24–27^. Thus, formation of each phase should be defined by a distinct saturation concentration ^24–28^. We denote saturation concentrations for fibril formation and condensation as *c*_sf_ and *c*_sc_, respectively. If *c*_sf_ < *c*_sc_, then condensates are metastable phases and fibrils are the thermodynamically stable ground state (**Figure 1A**). The free energy gap ΔΔ*G*_gap_ between condensates and fibrils defines the thermodynamic metastability of condensates with respect to fibrils. Free energy barriers define kinetic metastability (**Figure 1A**). For condensates to be crucibles for fibril formation, the thermodynamic requirement is that *c*_sf_ > *c*_sc_. If this were indeed true, then fibrils would represent a kinetic trap, rather than a thermodynamic ground state, and given enough time, fibrils would convert back to condensates with an input of energy.

**Figure 1:**
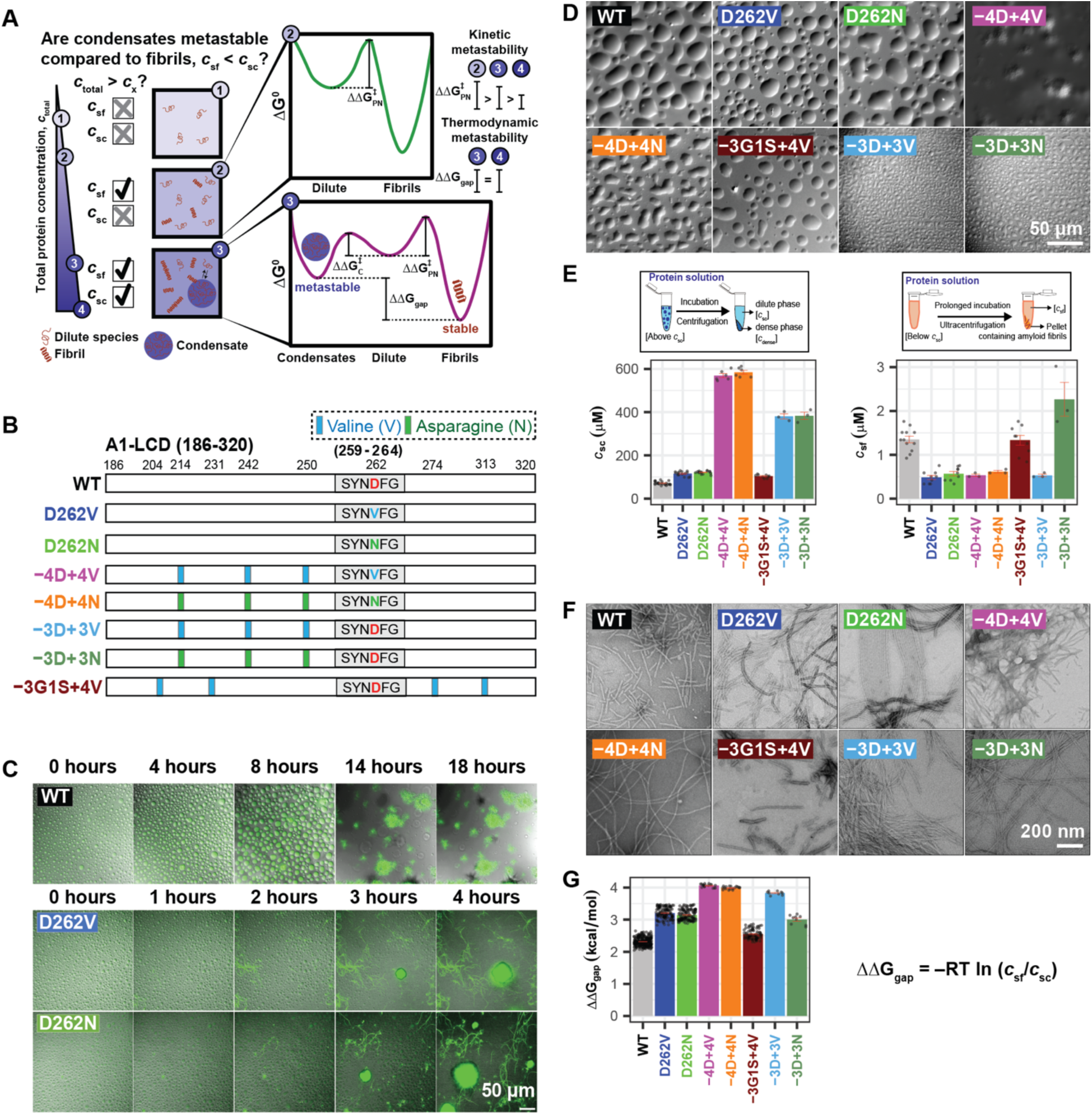
Pathogenic mutations reduce the metastability of A1-LCD condensates. **(A)** Schematic of the implications of measurable threshold concentrations *c*_sf_ and *c*_sc_ for the thermodynamic driving forces that govern condensate versus fibril formation. We depict expectations for the scenario where *c*_sf_ < *c*_sc_. If the total concentration, designated as *c*_total_, is lower than *c*_sf_ and *c*_sc_, then dispersed monomers are thermodynamically favored. If *c*_sf_ < *c*_total_ < *c*_sc_, then the thermodynamic driving forces favor separation of the system into two phases namely, dispersed monomers that coexist with fibrils. Kinetically, the barrier to nucleating fibrils will be governed by ΔΔG^‡^_PN_, the barrier to primary nucleation. For *c*_total_ > *c*_sc_, the metastable condensates serve as an off-pathway sink, and the barrier to nucleating condensates is set by ΔΔG^‡^_c_. The sink potential of condensates is quantified by ΔΔG_gap_ = –*RT*ln(*c*_sf_/*c*_sc_) which is the difference in standard state free energies of the fibril versus condensate phases. Here, *R* is the ideal gas constant, and *T* is the temperature of the system. **(B)** Schematic of the sequence architectures showing a steric zipper motif (S_259_YNDFG_264_, denoted as grey rectangle) within A1-LCD WT, pathogenic mutants, and designed variants that encompass the pathogenic mutations (−4D+4V/N and −3D+3V/N) or the WT sequence (−3G1S+4V). Vertical bars indicate the positions of substitutions to Val (light blue) or Asn (green). **(C)** Time course of superimposed differential interference contrast (DIC) and thioflavin T (ThT) fluorescence microscopy images of solutions of A1-LCD variants showing condensates and the appearance of fibrils. These panels were extracted from **Supplementary Videos 1-3**. **(D)** DIC micrographs of condensates of A1-LCD variants formed under conditions where *c*_total_ > *c*_sc_, which refers to the systems being supersaturated with respect to the condensate threshold. Data are shown for WT (*c*_total_ = 120 μM, *S*_c_ = 0.52), D262V (*c*_total_ = 200 μM, *S*_c_ = 0.55), D262N (*c*_total_ = 200 μM, *S*_c_ = 0.50), –4D+4V (*c*_total_ = 950 μM, *S*_c_ = 0.51), –4D+4N (*c*_total_ = 950 μM, *S*_c_ = 0.50), –3D+3V (*c*_total_ = 500 μM, *S*_c_ = 0.27), −3D+3N (*c*_total_ = 500 μM, *S*_c_ = 0.27), and –3G1S+4V (*c*_total_ = 180 μM, *S*_c_ = 0.54). **(E)** (Top left) Schematic depicting the method ^30^ used to measure *c*_sc_, and (bottom left) plot of *c*_sc_ of A1-LCD variants at 20°C, (top right) schematic depicting the method used to obtain *c*_sf_, and (bottom right) plot showing *c*_sf_ of A1-LCD variants obtained after 14 days of incubation of samples of 10 μM concentration at 20°C. Individual data points from replicate experiments are shown along with mean ± standard error in the estimate of the mean (SEM). **(F)** Negative-stain TEM images showing fibrils formed by A1-LCD variants in the pellets obtained after ultracentrifugation in the assay in (D, right). **(G)** Plot of ΔΔ*G*_gap_ for all A1-LCD variants. Individual values of ΔΔ*G*_gap_ from all possible pairs of *c*_sc_ and *c*_sf_ are shown along with the mean ± SEM. All experiments were performed in 40 mM HEPES buffer, pH 7.0 and 150 mM NaCl. Also see **Figure S1**.

Are condensates as a whole thermodynamically metastable or are fibrils kinetic traps? This rests on whether *c*_sf_ < *c*_sc_ or *c*_sf_ > *c*_sc_, and adjudication requires systematic measurements of *c*_sf_ and *c*_sc_. We find that *c*_sf_ < *c*_sc_ for all A1-LCD mutants and designed variants studied. Increased thermodynamic metastability of condensates increases the potential for condensate interiors to be sinks that suppress fibril formation. In accord with recent findings, we find that the interfaces of condensates serve as locations of heterogeneous nucleation. Together, our results highlight the dual roles of condensates and establish that tunable metastability of condensates modulates the rates of fibril formation.

## RESULTS

### Contributions of different residues to driving forces for condensate versus fibrils

We characterized the interplay between condensation and fibril formation for A1-LCD, which is the PLCD from hnRNPA1 (**Figure 1B**). The mutations D262N and D262V cause familial forms of ALS and MSP ^13^. *In vitro*, the A1-LCD variants (**Figure S1A**) were exchanged from a denaturing environment to a native buffer without excess salt. Addition of NaCl to a final concentration of 150 mM induced condensation (**Figure 1C, 1D**). The dense phases were fluid-like, undergoing fusion, and showing wetting behaviors. After several hours, fibrils were visible by light microscopy. Pathogenic mutations D262V and D262N enhanced fibril formation rates when compared to the WT A1-LCD (**Figure 1C**) ^13,29^.

Next, we measured *c*_sc_ and *c*_sf_ for different A1-LCD variants *in vitro*. We separated the dilute and dense phases using centrifugal sedimentation ^30,31^ and measured the concentrations of protein remaining in the dilute phase to quantify *c*_sc_ for each variant. The pathogenic mutations increased *c*_sc_, pointing to weakening of the driving forces for condensation (**Figures 1E** and **S1B**). We designed additional variants, mutating all four Asp residues to either Val or Asn (−4D+4V and −4D+4N, respectively), or mutating only three Asp to Val or Asn leaving the site of the pathogenic mutation (D262) intact (−3D+3V and −3D+3N, respectively). Increasing number of Asp substitutions results in a more positive net charge per residue (NCPR), and in accord with previous work this increased *c*_sc_ values compared to WT A1-LCD and the pathogenic mutants D262V and D262N (**Figures 1E** and **S1B, S1C**) ^31^.

Residues in A1-LCD and related PLCDs have been classified as stickers or spacers based on hierarchies of interactions mediated by different residues ^30–32^. Stickers form reversible physical crosslinks with one another, whereas spacers contribute directly to phase separation by determining solvation preferences, adding weak interactions, and affecting the cooperativity of inter-sticker interactions ^33^. To probe the effects of Val independently from the effects of increasing the NCPR, we replaced three Gly residues and one Ser residue with Val to generate the variant −3G1S+4V. This variant had a higher *c*_sc_ value relative to the WT A1-LCD, demonstrating that, despite their hydrophobicity and β-branched character, Val residues function as phase separation-weakening spacers, at least in the context of the A1-LCD (**Figures 1E** and **S1B**).

Next, we measured fibril stabilities by measuring *c*_sf_. Operationally, we defined *c*_sf_ as the concentration of protein that is left in solution after fibrillization is complete ^25,26,28,34^. After 14 days of incubation with shaking, the soluble protein concentration stabilized in the low micromolar range, independent of the input concentration (**Figures 1E** and **S1D**), and we observed fibrils in the pellet by transmission electron microscopy (TEM) (**Figure 1F**) and thioflavin T (ThT) fluorescence microscopy (**Figure S1E**). The *c*_sf_ values were lower than the corresponding *c*_sc_ values, showing that fibrils were globally stable, whereas condensates were metastable in comparison (**Figure 1A**). The pathogenic mutants had lower *c*_sf_ values than the WT A1-LCD, implying that they formed more stable fibrils. This is attributable to the substitution of D262 with V or N, which generates a steric zipper motif ^35^ and enhances the stability of fibrils ^13,36^ (**Figure S1F**). The *c*_sf_ values for the variants −4D+4V and −4D+4N were similarly low, showing that the effect of additional Asp substitutions on fibril stability is minimal. While the −3G1S+4V variant had a *c*_sf_ value that is equivalent to that of WT A1-LCD, the *c*_sf_ value for the −3D+3V variant resembled that of the pathogenic mutants although it lacks the steric zipper motif. This suggests that auxiliary zippers present in the sequence (**Figure S1F**) likely contribute to fibril stability. It appears that the effects of auxiliary zippers are masked in the −4D+4V/N variant because the strong steric zipper motif (^259^SYNDFG^262^) present at the site of the pathogenic mutation has a strong saturating effect on fibril stability. By contrast, the −3D+3N variant had the highest *c*_sf_ value of all variants (**Figure 1E**), suggesting that the interoperability of Asn with Val is context-dependent. Sequence-specific changes to *c*_sf_ are suggestive of the possibility that the −3D+3N variant likely forms fibrils of different polymorphs and stability, thus requiring a separate investigation of stability and high-resolution structural studies ^37^.

### A1-LCD condensates are metastable, and pathogenic mutations reduce this metastability

Our results show that pathogenic mutations within A1-LCD stabilize fibrils and destabilize condensates by lowering *c*_sf_ and increasing *c*_sc_ with respect to WT A1-LCD. This suggests that fibrils and condensates, which are ordered and disordered phases, respectively, are influenced differently by the pathogenic mutations. We quantified the metastability of condensates with respect to fibrils as the difference in standard state free energies of fibrils (Δ*G*°_f_) versus condensates (Δ*G*°_c_) (**Figure 1A**). Hence, ΔΔ*G*_gap_ = –*RT* ln(*c*_sf_ / *c*_sc_) where *R* = 1.98×10^-3^ kcal/mol-K is the ideal gas constant and *T* = 293 K. An increased free energy gap (ΔΔ*G*_gap_) implies decreased metastability of condensates compared to fibrils. If ΔΔ*G*_gap_ approaches zero, then condensates have increased metastability when compared to fibrils. We find that ΔΔ*G*_gap_ increases for pathogenic mutants (**Figure 1G**), showing that pathogenic mutations weaken condensate metastability. Additional mutations, including −4D+4V and −4D+4N, were akin to pathogenic mutations and had high values of ΔΔ*G*_gap_, implying low metastabilities. The −3D+3V variant has a ΔΔ*G*_gap_ between that of the pathogenic mutants and the −4D+4V/N variants. The *ΔΔG*_gap_ value for −3D+3N was between that of the WT and the pathogenic mutants (**Figure 1G**).

How do thermodynamically metastable condensates contribute to fibril formation? The answer comes from the Ostwald rule of stages ^38^: Kinetically accessible states are those that are closest in free energy to the starting state, and this need not be the globally stable phase. We therefore expected thermodynamically metastable condensates to be sinks that form more readily than fibrils under conditions where the total protein concentration is greater than *c*_sc_. Under these conditions, we indeed observed that condensates form more readily than fibrils (**Figure 1C**). This has been interpreted to mean that condensates are crucibles for fibril formation. However, such an interpretation implies that condensates are the stable states when compared to fibrils, which is not the case.

### Metastable condensates decelerate conversion to globally stable fibrils

The reduced metastability of condensates formed by pathogenic mutants suggests that the driving forces for fibril formation should be enhanced by these mutations. To test for this, we characterized the kinetics of fibril formation using the rate of gain of ThT fluorescence ^39^. These measurements were performed across a range of protein concentrations where the dispersed phase is subsaturated (*c*_sf_ < *c*_total_ < *c*_sc_) or supersaturated (*c*_total_ > *c*_sc_) with respect to condensates (**Figures 2A-2C**, and **S2** for repeats of the experiments).

**Figure 2:**
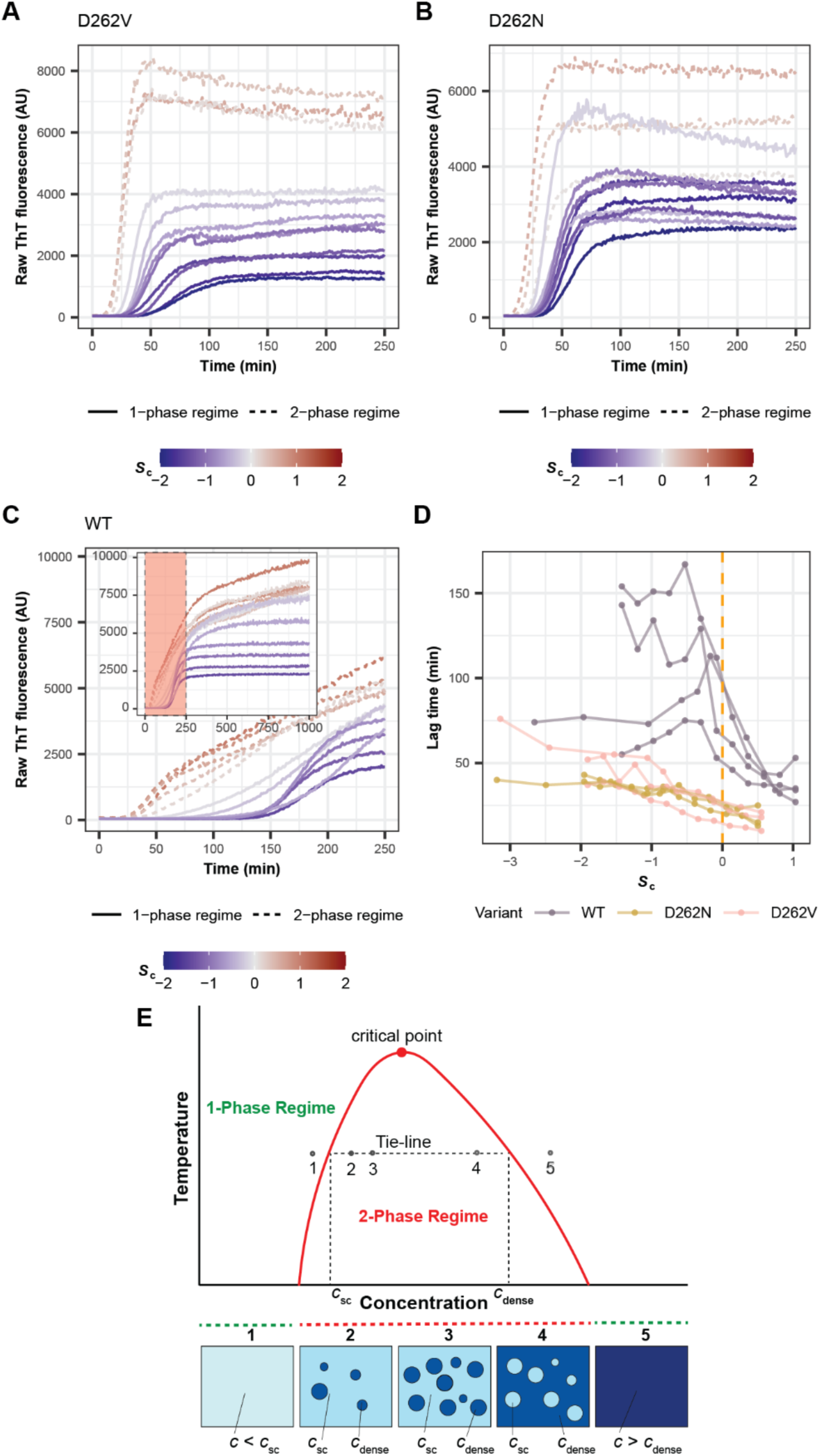
Metastable condensates decelerate conversion to globally stable fibrils. **(A, B, C)** Kinetics of fibril formation for A1-LCD variants (A) D262V, (B) D262N and (C) WT monitored by ThT fluorescence across a range of total protein concentrations (*c*_total_) that are subsaturated (*S*_c_ < 0, solid blue lines) or supersaturated (*S*_c_ > 0, dashed red lines) with respect to condensates. (C) Inset shows the complete time course. Data from a single representative experiment are shown; for replicate experiments see **Figure S2**. **(D)** Lag time of fibril formation (time at which ThT fluorescence reaches 10% of plateau value) for A1-LCD variants. The total protein concentrations are quantified as the degree of supersaturation relative to condensate formation (*S*_c_) and fibril formation (*S*_f_). Data from independent experiments using similarly prepared samples are represented by separate line plots. The dashed vertical line corresponds to *S*_c_ = 0. **(E)** Schematic of phase diagram of a UCST-type phase transition as observed for A1-LCD. The region below the coexistence curve (red) corresponds to the two-phase regime, where the sample contains condensates. The bottom panel is a visual representation of the sample under each condition. Notably, *c*_sc_ and *c*_dense_ are constant at increasing total protein concentration. Also see **Figures S3** and **S4.**

We plotted lag time values against the degree of supersaturation, *S*_c_ = ln(*c*_tot_ / *c*_sc_), of the dispersed phase with respect to condensates (**Figure 2D**). When the dispersed phase was subsaturated with respect to condensates, increases to protein concentration of the pathogenic mutants led to decreases in the lag times for fibril formation and increases in the overall rate of fibril growth (**Figures 2A,2B,2D** and **S2**). The rate of nucleation increases and the barrier to nucleation decreases as the dispersed phase becomes increasingly supersaturated with respect to fibrils ^40–43^. For concentrations that are above *c*_sc_, the dispersed phase is supersaturated with respect to both fibrils and condensates. Here, we observed further decreases in the lag time for fibril formation and increases in the slope that characterizes the growth of fibrils (**Figures 2D** and **S2**). Note that for *c*_total_ > *c*_sc_, condensates form, and protein concentrations in coexisting dilute and dense phases are fixed (**Figure 2E**). Only the fraction of molecules distributed across the phases and hence the volume occupied by the condensates should change. And yet we observed increased rates of initial fibril formation as *c*_total_ increased above *c*_sc_, as indicated by the decreasing lag times (**Figure 2D**). The kinetics of fibril formation of variants −4D+4V and −4D+4N, which exhibit even lower condensate metastabilities, had similar characteristics relative to those of the pathogenic mutants (**Figure S3**).

Increased nucleation rates with increase of *c*_total_ beyond *c*_sc_ cannot be explained by increased supersaturation with respect to *c*_sf_. This raised the question of how condensates might contribute to enhanced nucleation. Recent experiments have shown that condensate interfaces accelerate fibril formation ^20–22,44–48^. We used fluorescence microscopy with ThT as a marker to assess whether nucleation and initial growth originates at condensate interfaces. Low ThT fluorescence throughout condensates was observed at initial time points, in agreement with the expected small enhancement of fluorescence of molecular rotors by the high viscosity of the condensate interior ^49^ (**Figure S4**). Over time, ThT fluorescence increased at interfaces of condensates, suggesting that these regions promote fibril nucleation and initial growth. This effect is a plausible explanation for the observed decrease in lag time and increase in the rate of fibril formation with increasing volume fraction of condensates. This is in line with recent work showing that aging condensates of FUS proteins comprise cores and shells ^21^, suggestive of nucleation at the interface and preservation of soluble protein in the interior.

Unlike the pathogenic mutants, the condensates of WT A1-LCD have higher metastability, characterized by smaller values of ΔΔ*G*_gap_. For the WT A1-LCD, we found that fibril formation was characterized by longer lag times and slower growth rates (**Figures 2C, 2D** and **S2** for repeats). When *S*_c_ = 0 was crossed, the lag times continued to decrease monotonically, although more strongly for WT A1-LCD than for the pathogenic mutants (**Figure 2C, D**). However, the increased metastability of condensates led to distinct features in the ThT traces of WT A1-LCD (if *S*_c_ > 0) that are not expected for typical nucleation-and-growth models. Above *c*_sc_, fibril formation showed biphasic behavior, with an initial, relatively fast growth phase followed by a phase with reduced growth rate. The implication is that condensate interfaces enhance nucleation and initial fibril growth (**Figure S4**). However, the second phase with reduced growth rate is likely a reflection of metastable condensates serving as sinks for soluble proteins. We also observed similar biphasic kinetics for the −3G1S+4V variant in samples that were supersaturated with respect to condensates (**Figure S3**). Overall, there is an interplay between the accelerating effects of condensate interfaces and the sink potential, quantified in terms of *ΔΔG*_gap_, of condensate interiors. The latter seems to have a larger effect on fibril forming kinetics as ΔΔ*G*_gap_ decreases below ∼2.5-3.0 kcal/mol (**Figure 1G**).

Designed variants −3D+3V and −3D+3N, where we preserved the Asp residue in position 262, allowed us to test whether sequence-specific values of ΔΔ*G*_gap_ would determine the extent of suppression of fibril growth by condensates. Both variants had *c*_sc_ values between those of the WT and the −4D+4V/N variants. The ΔΔ*G*_gap_ value for the −3D+3V variant was between that of the pathogenic mutants and −4D+4V/N variants. In accordance with expectations, ThT traces for −3D+3V variant resembled those of the pathogenic mutants and −4D+4V/N variants (**Figure S3D**). By contrast, the −3D+3N variant with a ΔΔ*G*_gap_ value between that of the WT and the pathogenic mutants (**Figure 1G**) had biphasic ThT traces similar to the WT (**Figure S3E**). Therefore, sequence-specific values of ΔΔ*G*_gap_ determine the potential of condensates to suppress fibril growth.

### Modeling predicts that slow protein efflux from condensates slows fibril formation

Next, we analyzed the data for the kinetics of fibril formation using a numerical chemical kinetics approach ^50,51^. The model is parsimonious in its design (**Figure 3A**). It includes three species to mimic the dilute, dense, and fibrillar phases. Protein exchange between the dilute and dense phase species is reversible. Fibril formation is defined by rate constants for primary nucleation, elongation, and secondary nucleation ^52,53^. To minimize the number of fitting parameters, we assumed that fibril formation was irreversible on the timescale of the simulations. When *S*_c_ < 0, species corresponding to condensates are absent; accordingly, the influx and efflux rates were set to zero. When *S*_c_ > 0, the model includes an additional species corresponding to the dense phase.

**Figure 3:**
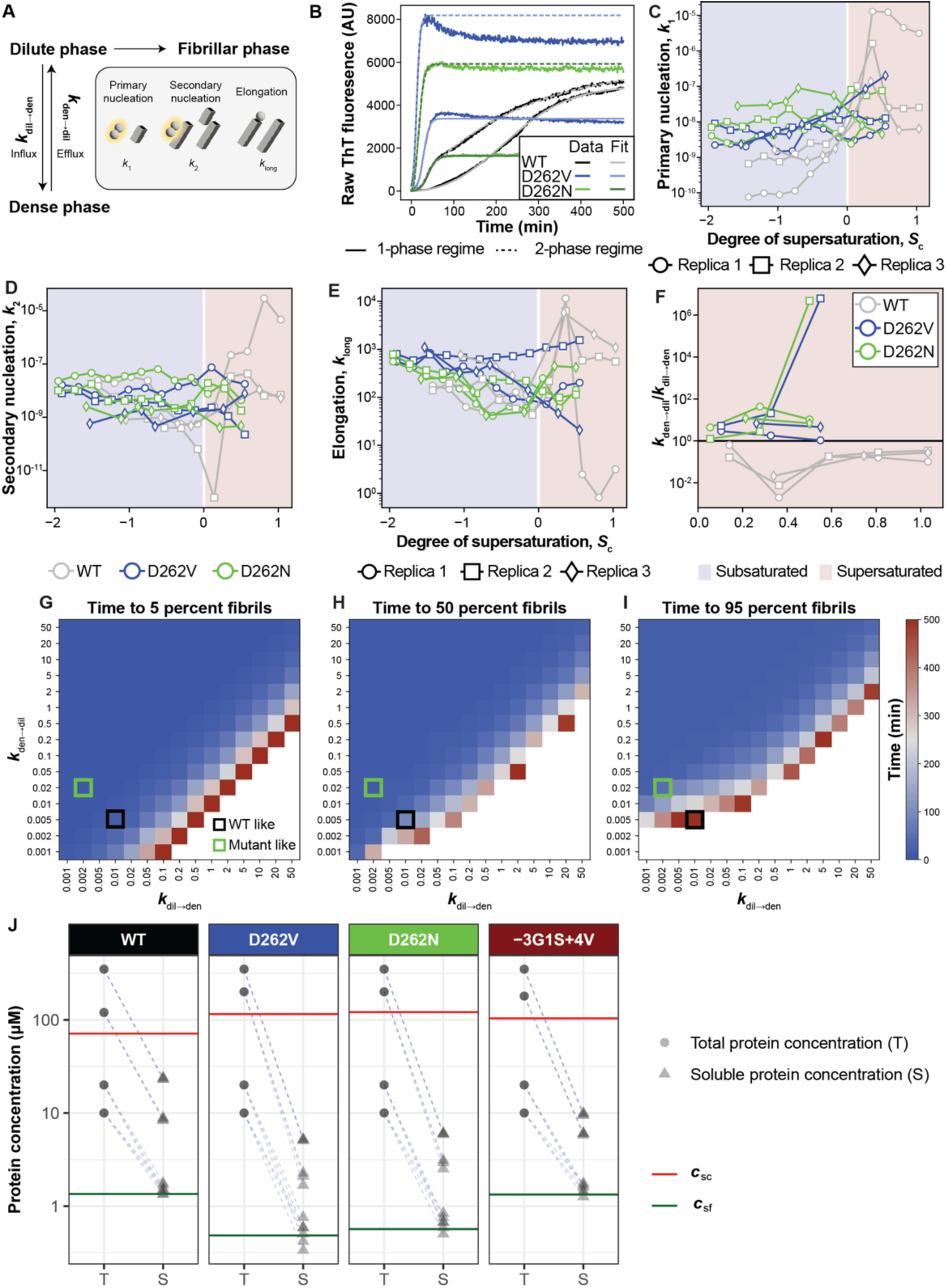
Slow protein efflux from condensates slows fibril formation. **(A)** Schematic of the chemical kinetics model used to fit ThT fluorescence curves (see Methods). **(B)** Examples of fits to ThT fluorescence traces for WT (*c*_total_ = 41 µM, *S*_c_ = –0.53), D262V (*c*_total_ = 65 µM, *S*_c_ = –0.57), and D262N (*c*_total_ = 65 µM, *S*_c_ = –0.61) in the absence of condensates (solid fit lines), and for WT (*c*_total_ = 128 µM, *S*_c_ = +0.59), D262V (*c*_total_ = 200 µM, *S*_c_ = +0.55), and D262N (*c*_total_ = 200 µM, *S*_c_ = +0.50) in the 2-phase regime (dashed fit lines). Concentrations were chosen to ensure similar degrees of subsaturation and supersaturation with respect to *c*_sc_ for each of the three constructs. **(C-F)** Parameters extracted from fitting ThT fluorescence traces to the chemical kinetics model for all reactions at total concentrations ≥ 17 µM and three replicas performed at sample ages of 0 days. Data were analyzed by fixing the supersaturation with respect to the condensation threshold. Thus, *S*_f_ will be system-specific. **(G-I)** The effect of titrating *k*_dil→den_ and *k*_den→dil_ on the time to reach **(G)** 5, **(H)** 50, and **(I)** 95 percent fibrils. Here, all other parameters are set to be constant with *k*_1_ = 1e-8, *k*_2_ = 5e-9, *k*_long_ = 500, and the total protein concentration, *c*_total_ = 200 µM. The total simulation time was 500 minutes. White values indicate that the target for percent of monomers incorporated into fibrils was not reached within this time. **(J)** Before- and-after plots comparing the input protein concentration and the concentration of soluble protein left in the supernatant after 14 days of incubation, when using different input concentrations above and below *c*_sc_. Data from individual measurements are shown. Red lines denote the *c*_sc_ values for each variant. When *c*_sf_ < *c*_total_ < *c*_sc_, the concentration in the supernatant consistently reaches the same value, which we interpret as *c*_sf_. (Data reproduced from Figures 1E and **S1D**.) However, when *c*_total_ > *c*_sc_, the amount of protein remaining in the supernatant increases, depending on the extent of supersaturation, pointing to the residual sequestration of soluble proteins in condensates even after 14 days of incubation and ultracentrifugation. Also see **Figure S5**.

We used the model to fit ThT traces for pathogenic mutants and WT for *S*_c_ < 0 and *S*_c_ > 0 (**Figure 3B**). From the fits to the model, we propose that the data are consistent with a mechanism whereby fibril formation of the WT is slower than that of pathogenic mutants because of slower primary nucleation for *S*_c_ < 0 (**Figures 3C-3E**). In the chemical kinetics approximation, secondary nucleation implicitly includes the contribution of condensate interfaces. For *S*_c_ > 0, the rates of primary and secondary nucleation increase for the WT beyond that of the mutants while the rates for the mutants stay relatively unchanged. Thus, dense phases have smaller effects on pathogenic mutant fibril formation rates. The model also predicts that the ratio of efflux to influx rate constants should be lower for the WT relative to the pathogenic mutants (**Figure 3F**). We observed a rapid decay of the fraction of protein in the dilute and dense phases for the pathogenic mutants, that coincides with rapid fibril growth (**Figure S5**). In contrast, the protein fraction in the dense phase decayed more slowly for WT, leading to slower fibrillar growth (**Figure S5**).

Next, we performed a parameter sweep to test how rate constants for influx and efflux influence the time it takes for 5%, 50% or 95% of the protein to convert to fibrils (**Figures 3G-3I**). All other rate constants were kept constant. A large range of ratios of *k*_dil→den_/*k*_den→dil_ were found to result in rapid incorporation of 5% of the protein (*t*_0.05_) into fibrils. However, above a certain ratio of *k*_dil→den_/*k*_den→dil_, we found that *t*_0.05_ rises sharply and fibril formation is suppressed, indicating that proteins are sequestered in dense phases when influx into condensates is much faster than efflux. Similarly, the time taken to incorporate 50% of protein into fibrils (*t*_0.5_) was found to be strongly influenced by the ratio of *k*_dil→den_/*k*_den→dil_. In this case, we also observed an influence of the absolute value of *k*_den→dil_, whereby below cutoff values of *k*_den→dil_, 50% incorporation into fibrils could not be reached within 500 minutes. This cutoff value was found to depend on the rates for nucleation and growth. The time to reach 95% incorporation into fibrils (*t*_0.95_) depends even more strongly on *k*_den→dil_ in addition to the ratio of *k*_dil→den_/*k*_den→dil_. Overall, the analysis shows that the exchange of material between species mimicking dilute phases and condensates, particularly the efflux from condensates, becomes limiting for fibril growth in the dilute phase. This is highlighted by the WT-like ratio of *k*_dil→den_/*k*_den→dil_ (**Figure 3I**, black square) corresponding to a longer time to reach 95% incorporation into fibrils compared to the pathogenic mutant-like ratio of *k*_dil→den_/*k*_den→dil_ (**Figure 3I**, green square). Overall, the chemical kinetics modeling suggests that species corresponding to dense phases should shrink over time. We tested this prediction using video microscopy, which showed condensates shrinking over time, and weakly metastable condensates were found to shrink more rapidly (**Videos S1-S3**). Taken together, the model predicts and the data show that pathogenic mutations weaken the sink potential of condensates, thereby tilting the balance in favor of fibrils.

To test the prediction that condensates are sinks for soluble protein, we asked if the presence of condensates slowed equilibration to the globally stable fibrillar state. We repeated measurements of the soluble protein concentration after two weeks of incubation with shaking, and the total protein concentrations exceeding *c*_sc_. (**Figure 3J**). Unlike the measurements performed when *c*_total_ < *c*_sc_, input concentrations above *c*_sc_ resulted in soluble protein concentrations above *c*_sf_ even after two weeks of incubation; the higher the input concentrations, the more soluble protein remained. These results suggest that the presence of condensates slows equilibration because metastable condensates act as sinks for soluble protein.

Taken together, while fibrils can be nucleated by condensate interfaces, the interiors of condensates function as sinks, and fibrils grow in dilute phases. We observed that pathogenic mutants formed fibrils that grow rapidly in a small number of places, with concomitant dissolution of condensates (**Videos S1, S2**, see also **Figure 1C** for stills). WT fibrils, in contrast, formed at condensate interfaces and grew slowly, without individual fibrils dominating fibril growth. WT condensates also shrink slowly, limiting WT fibril growth (**Video S3**, see also **Figure 1C** for stills).

### Enhancing the metastability of mutant condensates suppresses fibril formation

Condensates can be stabilized by introducing stronger stickers into A1-LCD ^31^. Replacing Phe and Tyr with Trp residues enhances the driving force for condensate formation ^54^. We asked if the pathogenic phenotype of D262V/N A1-LCD might be rescued by enhancing condensate metastability through the introduction of stronger stickers. We designed variants in which we mutated all or a subset of Phe and Tyr residues to Trp while keeping the sequence around the site of the disease mutation intact, resulting in allW, allW D262V, allW D262N, 11W D262V and 5W D262V variants (**Figure 4A**). These variants readily formed condensates (**Figures 4B** and **S6A,S6B**), and their *c*_sc_ values were lower than those of the parent variants (**Figure 4C**). However, the *c*_sf_ value for allW was equivalent to that of WT, and those for allW D262V and allW D262N were between those of the original pathogenic mutants and WT (**Figure 4D**). All five Trp variants formed fibrils within 14 days (**Figure 4E**). The 1Δ1Δ*G*_gap_ values for the new variants were smaller than those of WT and the original pathogenic mutants (**Figure 4F**), implying increased metastability of condensates when Phe and Tyr stickers were replaced with Trp.

**Figure 4:**
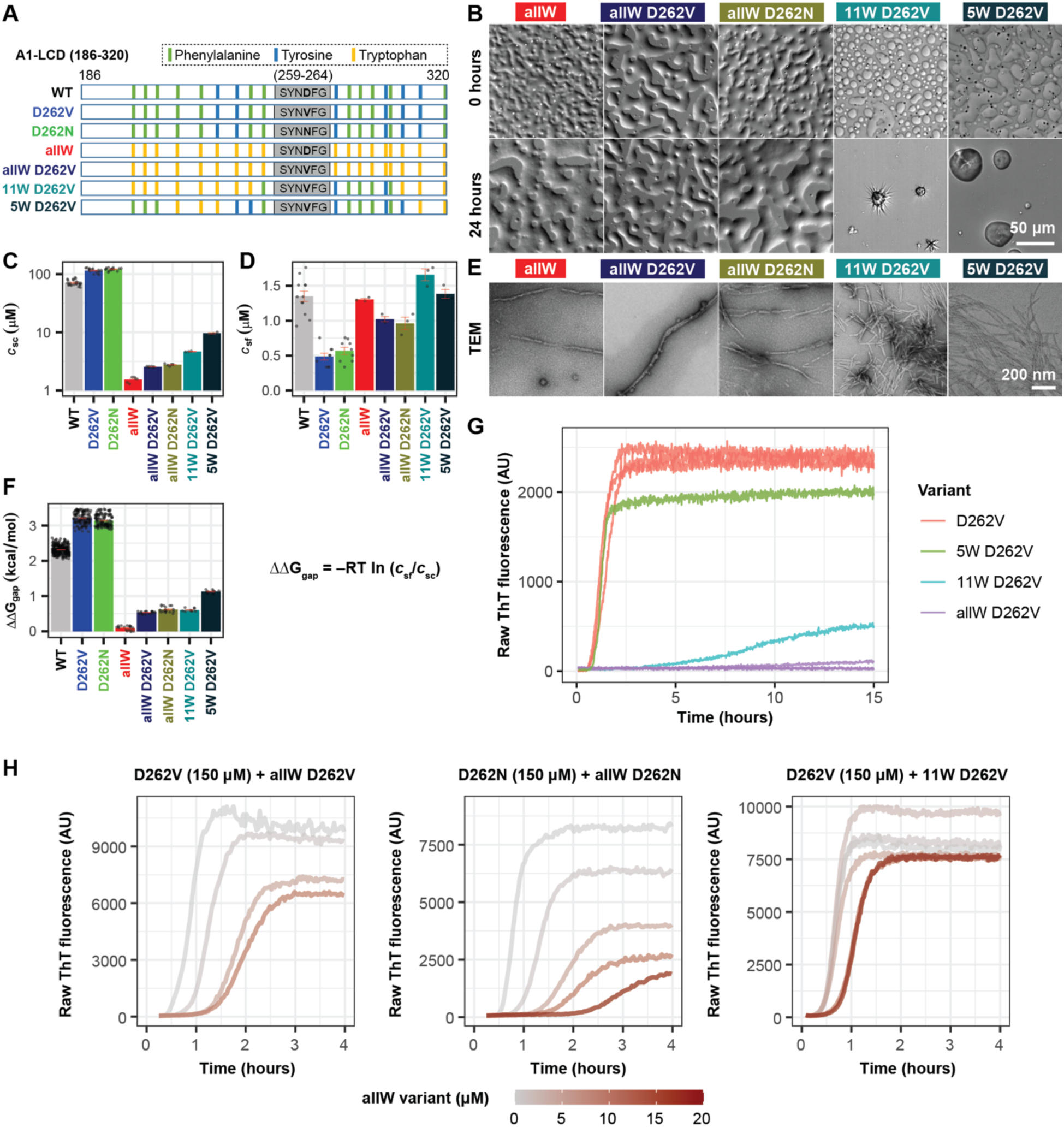
Enhancing the metastability of mutant condensates suppresses fibril formation. **(A)** Schematic of sequence architectures of A1-LCD wild-type, pathogenic mutants, and the Trp variants depicting the sites of mutations. Vertical bars indicate the position of Phe (green), Tyr (blue) or Trp (yellow) residues in the sequence. The motif surrounding the pathogenic mutation site (S_259_YNDFG_264_) is shown as a grey rectangle. **(B)** DIC images of condensates formed by the Trp variants at 0 hour and 24 hours of incubation. The structures formed by variants 11W D262V and 5W D262V after 24 hours are bundles of fibrils ^50^. **(C)** Measured *c*_sc_ values for WT A1-LCD, pathogenic mutants and Trp variants. **(D)** Measured *c*_sf_ values for these variants at 20°C. Individual measurements are shown along with mean ± SEM. Values for WT A1-LCD, D262V and D262N are taken from Figure 1E. **(E)** Negative-stain TEM images showing fibrils (and some oligomers) formed by Trp variants in the pellets obtained after ultracentrifugation in the assay in panel (C). **(F)** ΔΔ*G*_gap_ values for A1-LCD variants. Individual data points calculated from all possible pairs of replicate experiments in (C, D) are shown along with the mean ± SEM. **(G)** ThT fluorescence-monitored fibrillization kinetics of A1-LCD variants D262V, allW D262V, 11W D262V and 5W D262V at equal concentrations of 40 μM showing raw ThT fluorescence as a function of time. For additional concentrations see **Figure S6E**. **(H)** Suppression of fibril formation of the D262V disease mutant in the presence of increasing concentration of the allW D262V and 11W D262V variants, and of the D262N disease mutant in the presence of increasing concentration of the allW D262N variant. All experiments were carried out in 40 mM HEPES buffer, pH 7.0 and 150 mM NaCl. Also see **Figure S6**.

ThT assays confirmed a strong kinetic suppression of fibril formation for variants allW D262V and 11W D262V but less so for 5W D262V (**Figure 4G**). Thus, the effect from incorporating Trp residues was titratable. To test if inhibition worked in *trans*, we added increasing concentrations of allW D262V/N or 11W D262V variants to a constant concentration of pathogenic mutant protein D262V/N; the resulting condensates concentrated both proteins (**Figures S6D,E**). Fibril formation was delayed and reduced with increasing concentrations of the Trp variants (**Figure 4H**). Thus, condensates with higher metastability suppress or slow down fibril formation.

### Trp variants slow both fibril nucleation and growth

Given the effects of the allW and 11W variants on lag time and fibril growth of the pathogenic mutants (**Figure 4H**), we considered two factors in addition to the sink potential of condensates as possible mechanisms for the suppression of fibril formation. We asked if fibril formation would be delayed by the presence of Trp variants in the absence of condensates. To test for this, we added increasing concentrations of Trp variants to a fixed concentration of pathogenic mutant D262V, while maintaining concentrations in the regime of *S*_c_ < 0 (**Figure 5B**). The presence of variants 5W D262V and 11W D262V did not suppress fibril formation of pathogenic mutant protein D262V, but the addition of allW D262V delayed fibril formation modestly. The allW variant likely forms clusters when *S*_c_ < 0 ^55^ that potentially influence fibril formation in subsaturated solutions.

**Figure 5:**
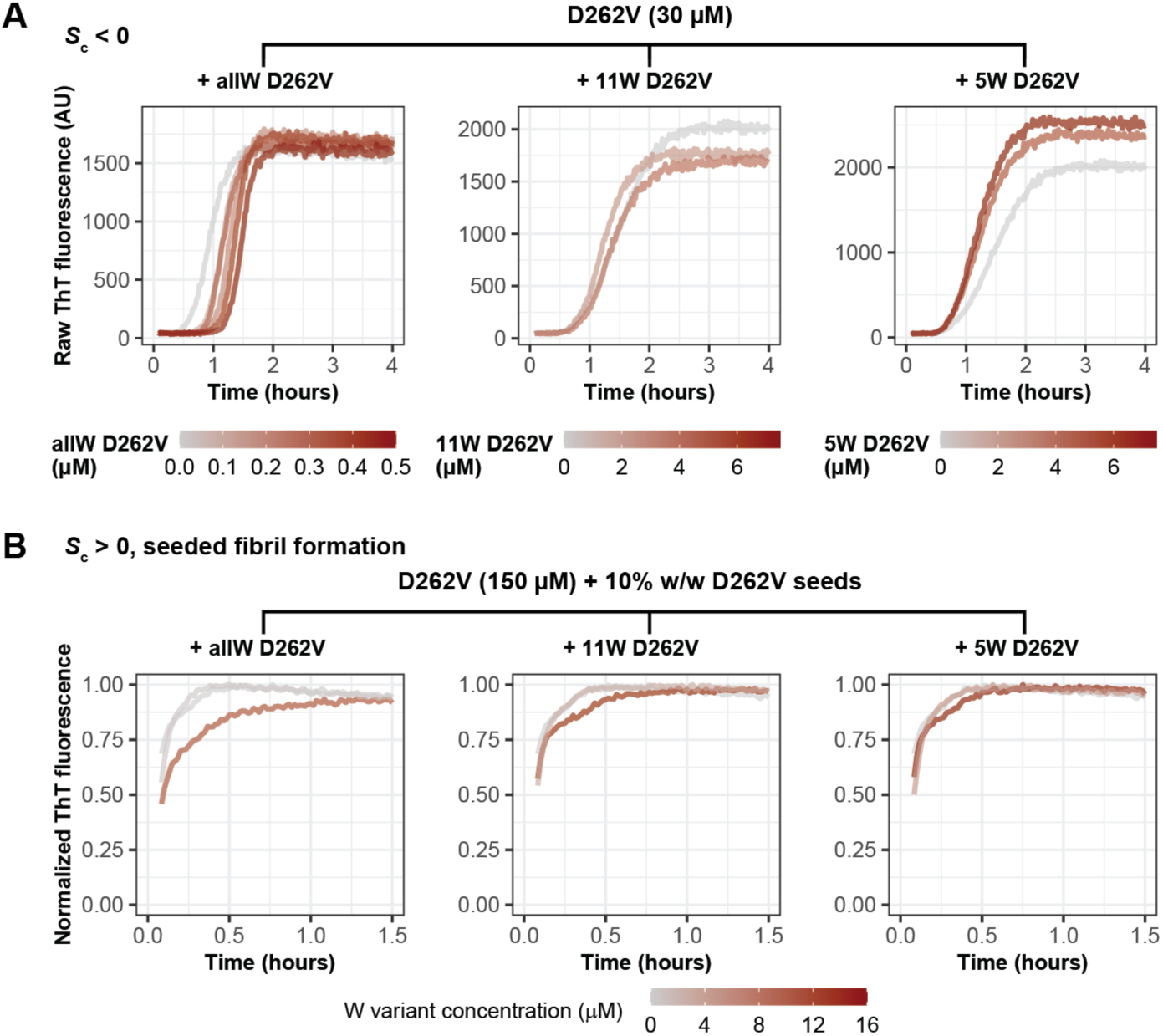
Trp mutants slow the nucleation and growth of fibrils. **(A**) Effect of Trp variants (allW D262V, 11W D262V, 5W D262V) on the fibril formation kinetics of the pathogenic mutant of A1-LCD in the absence of condensates. **(B)** Effect of Trp variants on the kinetics of seeded aggregation (in presence of 10% w/w fibril seeds of A1-LCD D262V). To compare lag time and growth rates, the ThT fluorescence data are shown normalized to the plateau value.

Next, we considered the effect of Trp variants on nucleation of fibrils at condensate interfaces. We performed seeded fibrillization kinetics experiments using the pathogenic mutant protein D262V in the presence of D262V fibril seeds and increasing concentrations of Trp variants (**Figure 5C**). This was done to separate contributions from suppressing nucleation versus fibril growth. Under these conditions, the presence of Trp variants reduced the slope of ThT traces, and the size of the effect was titratable by the concentrations of Trp variants and extent of Trp substitutions. Overall, condensates of higher metastability suppress fibril growth. In addition, condensates with particularly high metastability delay nucleation at condensate interfaces compared to condensates with lower metastability (**Figures 4G, 4H** and **Figures S6A, S6B**). A plausible explanation comes from recent experimental observations ^56^ and computations ^56–58^. These studies show that enhancing the driving forces for phase separation lead to more parallel orientations of scaffold molecules at interfaces, and we reason that this can affect the probability of nucleation at interfaces ^56,57^.

### Metastability of condensates governs the timescales of their dissolution

Chemical kinetics modeling suggested that the efflux of soluble protein from the dense to the dilute phase can become rate-limiting. To test whether metastability governs the efflux rates, we performed measurements of the kinetics of condensate dissolution. Single condensates were optically trapped in a microfluidic flow chamber and moved to a flow channel that contained only buffer, effectively jumping the system to strongly sub-saturating solutions that promote dissolution (**Figure 6A, Videos S4-S6**). The dissolution of condensates was followed by measuring their cross-sectional area over time. Flow in the channel accelerates condensate dissolution compared to the absence of flow. Accordingly, we compared different variants relative to each other instead of interpreting absolute dissolution rates.

**Figure 6:**
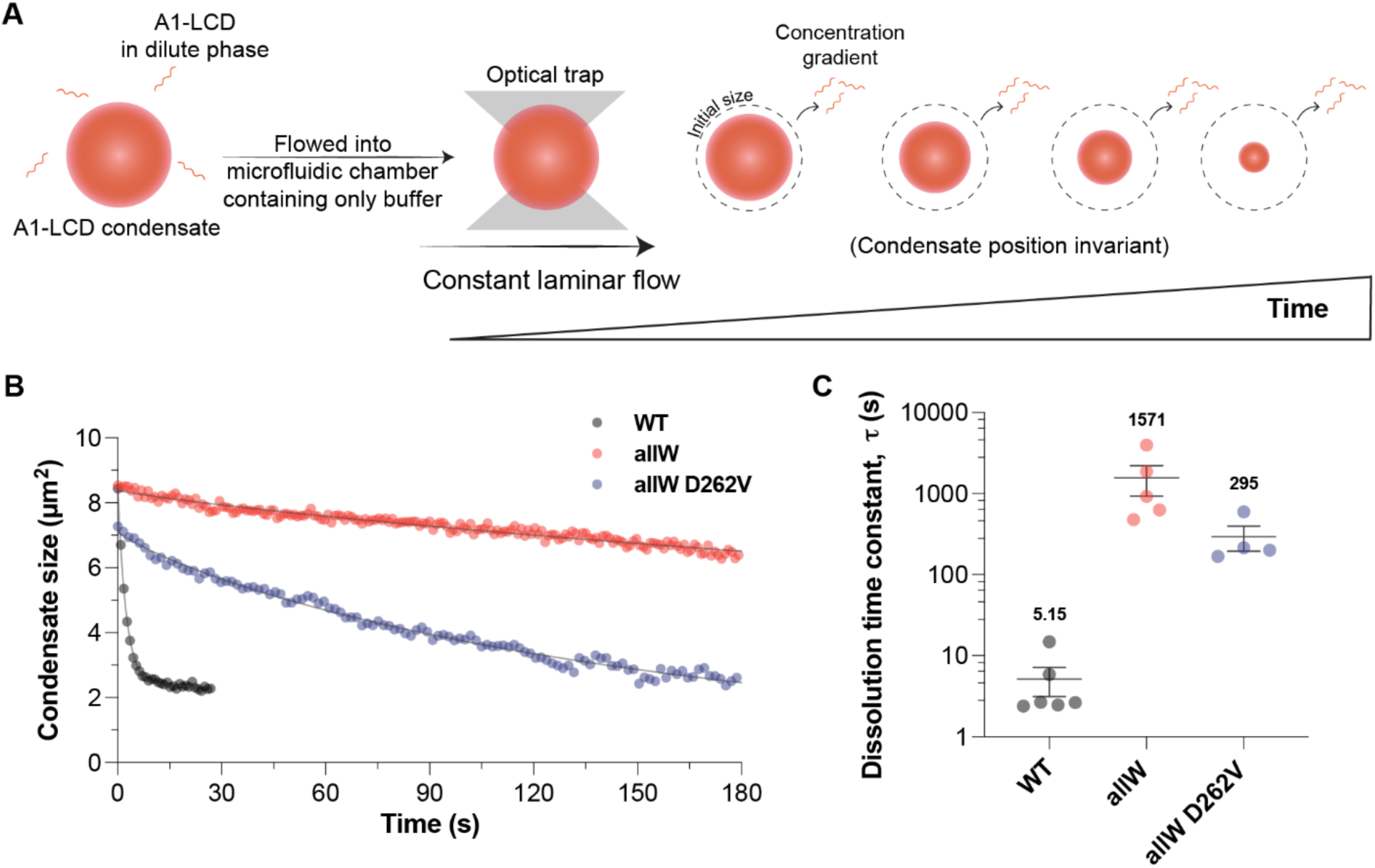
Condensate metastability governs the kinetics of protein efflux. **(A**) Schematic of the experimental setup for condensate dissolution measurements using an optical tweezer and a laminar flow microfluidic cell. **(B)** Representative decay traces of condensate size as a function of time for different A1-LCD variants, fitted using a stretched exponential function. See also **Videos S4-S6**. **(C)** Dissolution time constants of A1-LCD variants extracted from decay traces of individual condensates. Data from individual measurements are shown along with mean ± SEM. The mean dissolution time constant (in sec) of each variant is indicated. All experiments were performed in 40 mM HEPES buffer, pH 7.0, and 150 mM NaCl and the concentration of each A1-LCD variant was as follows: WT, 300 μM; AllW, 100 μM; AllW D262V, 200 μM.

Recent work has shown that condensates are viscoelastic materials ^54^ defined by a spectrum of relaxation times ^59^. Dissolution dynamics of such systems can be modeled using the Kohlrausch-Watt-Williams (KWW) function ^60,61^. We fit each of the dissolution traces to a KWW stretched exponential defined by the fastest relaxation time ι− and an exponent β that quantifies the ruggedness of the free energy landscape that gives rise to a distribution of barriers.

WT condensates dissolved within a few seconds under flow with a time constant τ of 5.15 s (**Figures 6B, 6C)**. The dissolution dynamics followed a single exponential decay with the exponent β being close to 1 (see **Methods**). Condensates formed by the pathogenic mutant protein D262V dissolved rapidly. The protein also aggregated in the microfluidic channel, and we were thus unable to capture its condensate dissolution kinetics. To better explore the effect of pathogenic mutations, we compared the dissolution kinetics of condensates formed by the allW and allW D262V variants, which had time constants τ of 1571 s and 295 s and exponents β of 0.84 and 0.85, respectively (**Figures 6B, 6C, Videos S5, S6**). These results suggest the following: The slower dissolution of Trp variants demonstrates that mutations that stabilize the percolated networks within condensates ^32,54,59,62^ will slow the efflux of soluble protein from dilute phases. The values of β that are lower than 1 suggest the presence of a distribution of barriers that slows down protein efflux from the percolated network within condensates ^32,54,59^. Pathogenic mutations destabilize condensates and increase the efflux rates of soluble protein. Overall, these results suggest that highly metastable condensates can suppress fibril growth through slow efflux of protein from condensates.

### Mutations that increase the metastability of A1-LCD condensates rescue a SG disassembly phenotype in cells

Finally, we asked if mutations that increase the metastability of condensates formed by homotypic interactions *in vitro* could also rescue a SG-related disease phenotype. The expression of pathogenic hnRNPA1 mutants slows energy-dependent SG disassembly dynamics ^63,64^. Delayed SG disassembly is a pathogenic phenotype in cultured cells ^14,63^, and we asked if this might be rescued by stabilizing PLCD-mediated condensate formation of hnRNPA1. We replaced the aromatic residues in the PLCD of a full-length hnRNPA1 construct with Trp while leaving the steric zipper motif and nuclear localization sequence unchanged. U2OS cells were transiently transfected with a construct proportionally expressing C-terminally FLAG-tagged hnRNPA1 variants and eGFP as a separate polypeptide via an internal ribosome entry site (IRES). The cells were subjected to heat stress for 1 hour at 43°C, which resulted in the assembly of SGs identified by poly-A binding protein (PABP) immunostaining. C-terminally FLAG-tagged hnRNPA1 variants colocalized with the SGs (**Figure S7A**).

Next, we transfected U2OS cells expressing the SG hub G3BP1-tdTomato from its endogenous locus ^1^ with these constructs (**Figure 7A**). This allowed for the tracking of SGs in live cells. After one day, the cells were subjected to heat stress for 1 hour at 43°C, which resulted in the assembly of G3BP1-positive SGs. The expression levels of hnRNPA1 variants and the extent of SG assembly in cells expressing the different hnRNPA1 variants were comparable (**Figures S7B,S7C**). Upon release of the stress, achieved by restoring the temperature to 37°C, the SGs disassembled over time (**Figure 7B** and **S7D-S7F**), and the rate of disassembly was quantified for cells with moderate expression of eGFP (**Figure 7C**).

**Figure 7:**
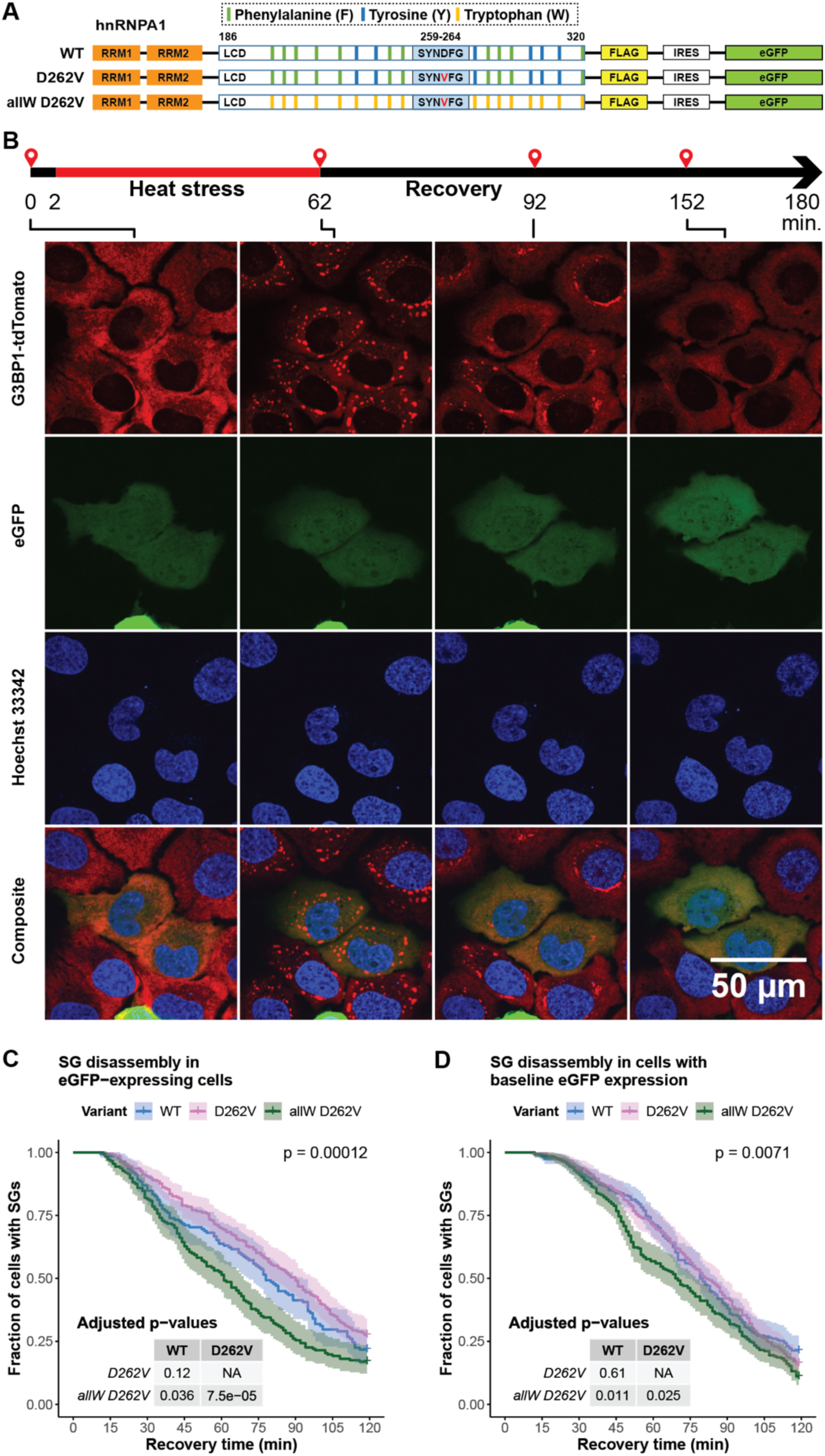
Mutations that increase the metastability of A1-LCD condensates rescue a SG disassembly phenotype in cells. **(A)** Schematic of the architecture of full-length hnRNPA1 variants with C-terminal FLAG tag and proportionately expressing eGFP via an interim IRES sequence, showing the location and identity of the aromatic residues in the LCD. **(B)** Representative micrographs of U2OS cells expressing the SG marker G3BP1-tdTomato (red) from its endogenous locus, that were transiently transfected with the WT hnRNPA1-FLAG-IRES-eGFP construct and subjected to 1 hour of heat stress at 43 °C, during which SGs become visible as punctate G3BP1 foci in the cytoplasm. In the recovery phase at 37 °C, the SGs dissolve over time. eGFP (green) identifies the cells expressing protein from the transfected construct. The cell nuclei are stained with Hoechst 33342 (blue). **(C)** Quantification of SG disassembly kinetics in cells expressing eGFP. The mean of three independent replicate experiments is shown. **(D)** Quantification of SG disassembly kinetics from the same experiments as in (C) but of cells without eGFP signal. (C,D) The mean of three independent replicate experiments is shown. The shaded regions represent the 95% confidence interval. The three variants were compared using a log rank test, and the overall p-value that tests for differences between variants is shown in the top right corner. For pairwise comparisons of variants, adjusted p-values (using the Benjamini-Hochberg correction) are shown in the table inset. Also see **Figure S7**.

Cells expressing the pathogenic hnRNPA1 mutant D262V showed delayed SG disassembly compared to cells expressing hnRNPA1 WT (**Figure 7C**). The effect of the mutation was comparable to previous observations ^63,64^. In contrast, in cells expressing the designed hnRNPA1 variant protein allW D262V, the SGs started to disassemble on timescales that were akin to those of cells expressing the WT protein and completed disassembly even faster than WT. We also analyzed SG disassembly in a population of cells that expressed zero to very low levels of hnRNPA1 variants as indicated by eGFP levels at background levels as a control. SG disassembly dynamics in these cells were identical within error for WT and D262V variants; transfection with the allW D262V variant seemed to confer a slight advantage (**Figure 7D**). This may be attributable to some cells expressing low levels of hnRNPA1 despite background levels of eGFP, attributable to relatively low levels of eGFP expression from the IRES (**Figure S7B**). Indeed, examples of cells expressing WT or D262V hnRNPA1 without obvious eGFP signal are detectable in our dataset (see **Figure S7A** for examples). Thus, to test the phenotypic effect of expression of each hnRNPA1 variant, we systematically compared SG disassembly dynamics for cells expressing and not expressing each of the constructs, as inferred from the presence or absence of eGFP fluorescence. While the expression of D262V delayed SG disassembly compared to cells that were transfected with the same construct but did not show measurable expression, the expression of WT and allW D262V variants conferred a disassembly enhancement (**Figure S7D-S7F**). These experiments suggest that even very low expression levels of hnRNPA1 variants influenced SG disassembly dynamics. Our results suggest that pathogenic mutants of hnRNPA1 with decreased metastability slow SG disassembly. Strikingly, this effect can be rescued by mutations that increase the metastability of condensates, leading to a wild type-like SG disassembly phenotype in cultured cells.

## DISCUSSION

Overall, our findings may be summarized as follows: Condensates are metastable relative to globally stable fibrils (**Figure 1A,1G**). Condensates form more readily than fibrils (**Figure 1C**) on sub-second timescales for A1-LCD ^30^, but the system converts to globally stable fibrils over the course of hours to days (**Figures 1C, 2** and **3J**). The metastability of condensates relative to fibrils is an important factor in determining the timescale of conversion to fibrils. Conversion can be slow if condensates have increased metastability defined by lower values of ΔΔ*G*_gap_ because condensates can serve as effective sinks for soluble proteins (**Figure 3J**) to limit the rate of efflux (**Figure 6**). The pathogenic mutations in the A1-LCD weaken the metastability of condensates to stabilize fibrils (**Figure 1E,1G**). Condensates formed by pathogenic mutants do not sequester soluble proteins as readily as WT condensates (**Figures 2** and **3**). Interfaces of condensates can nucleate fibril formation (**Figures 2** and **S4**), and mutations that affect condensate metastability also change the effectiveness of nucleation at interfaces (**Figure 5**). Fibril growth occurs in the dilute phase and is determined by the protein concentration in the dilute phase, which is kinetically regulated by the efflux of proteins from condensates (**Figures 3** and **6**). Introducing mutations that weaken the metastability of A1-LCD condensates also has a deleterious effect on SG disassembly in cells. Mutations that enhance the metastability of the condensates rescue the SG disassembly phenotype in cells even if they feature pathogenic mutations (**Figure 7C**). Our findings point to separable contributions of stickers that contribute to stabilization of disordered condensates and steric zippers that stabilize ordered, fibrillar phases ^63^. Thus, pathogenic mutations in A1-LCD appear to be separation-of-function mutations that destabilize condensates and stabilize fibrils.

We find that sticker-sticker interactions are important for preventing transformation into fibrils. We propose that in SGs, the network of heterotypic sticker-mediated interactions slow the formation of productive interactions of fibril-promoting motifs, thus suppressing fibril formation in the dense phase. This proposal is in agreement with the heterotypic buffering model put forward previously, in which heterotypic interactions in complex condensates suppress the conversion into fibrils formed by homotypic interactions ^65^. Recent work has suggested that the formation of separate TDP-43 condensates or demixing of TDP-43 within condensates, either via oxidative stress, protein unfolding stress, or optogenetic engineering, which cause loss of heterotypic buffering, promotes conversion to solid states and pathogenic processes ^19,66,67^. Kinetic stabilization against fibrillization has also recently been proposed for translation-repressing condensates formed by CPEB4 ^68^.

Given the tendency of condensate interfaces to nucleate fibril formation ^20–22,44,45^, mechanisms to passivate those interfaces may have evolved. These can include emulsification of condensates by specific proteins that accumulate at the interface ^69^. Prolonged stresses or the presence of destabilizing pathogenic mutations may render the buffering functions of condensates to be less effective.

### Limitations of the study

Our studies were performed using a single PLCD and variants thereof. Future work will focus on characterizing the effects of compositionally complex facsimiles of SGs. While our findings are likely generalizable at least for PLCD-containing systems, they may not apply to proteins with non-overlapping sticker and zipper sequence grammars. Additional work is needed to test whether phenotypes generated by the expression of pathogenic RBP mutants are rescued by stabilizing the PLD network in SGs. Novel methods will also be needed to follow the temporal evolution of all the relevant species that go beyond fibrils.

## ACKNOWLEDGMENTS

Microscopy images were acquired at the Cell and Tissue Imaging Center at SJCRH, which is supported by SJCRH and NCI (grant P30 CA021765). We thank Hong Joo Kim and Brian Freibaum, Richard Kriwacki, and Kresten Lindorff-Larsen for insightful discussions. This research was supported by grants from the National Institutes of Health R01NS121114 (to RVP and TM), grant R35NS097974 (to JPT), and grant R35GM138186 (to PRB), the St. Jude Research Collaborative on the Biology and Biophysics of RNP granules (to JPT, PRB, RVP, and TM), the American Lebanese Syrian Associated Charities (to JPT and TM), and the Air Force Office of Scientific Research (grant FA9550-20-1-0241 to RVP).

## AUTHOR CONTRIBUTIONS

Conceptualization: RVP, TM; Methodology: TD, FZ, MF, KMR, TSM, AS, JM, JPT, PRB, RVP, TM; Investigation: TD, FZ, MF, KMR, TSM, AS; Visualization: TD, FZ, MF, KMR, TSM, PRB, RVP, TM; Funding acquisition: JPT, PRB, RVP, TM; Supervision: PRB, RVP, TM; Writing – original draft: TD, FZ, RVP, TM; Writing – review & editing: all authors.

## SUPPLEMENTAL INFORMATION

Document S1, Figures S1 to S7, Tables S1 to S2.

Videos S1 to S6

## Methods

### Constructs

All constructs used for in vitro experiments are variants of the LCD of heterogeneous nuclear ribonucleoprotein 1 (hnRNPA1), and the coding sequences were either derived from the original A1-LCD WT plasmid (sequence not optimized for bacterial expression) ^30^ by site-directed mutagenesis, or synthesized by Thermo Fisher Scientific (codon-optimized for bacterial expression) with a sequence coding for an N-terminal TEV cleavage site (ENLYFQGS) and 5’ and 3’ attB sequences for Gateway cloning. Synthetic constructs were cloned into the expression vector (pDEST17 vector) through an LR reaction and expressed as fusion proteins containing a 6X His-tag and the TEV protease cleavage site. Following purification, cleavage with TEV protease was performed, after which the residues ‘GS’ remain at the N-terminus along with the sequence for A1-LCD (186-320). All sequences are shown in **Table S1** (bacterial expression constructs) and **Table S2** (mammalian expression constructs).

### Protein expression and purification

The non-codon optimized constructs of A1-LCD (WT, D262V, D262N) were expressed in *E. coli* BL21 (DE3) RIPL cells. All codon-optimized variants were expressed in *E. coli* BL21 (DE3) Gold cells. Cultures were grown in ZYM5052 auto induction media ^70^ at 37°C for 18-24 hours. Cell pellets were resuspended in lysis buffer (50 mM MES pH 6.0, 500 mM NaCl, 20 mM 2-mercaptoethanol), lysis was performed via sonication and the samples were centrifuged (30,000 g, 30 min) to pellet inclusion bodies containing the expressed protein. The pellet was further resuspended in 6 M guanidinium chloride, 20 mM Tris pH 7.5, 15 mM imidazole and solubilized overnight at 4°C with constant stirring. The solubilized pellet was centrifuged (30,000 g, 30-40 min) and the clear supernatant was loaded onto self-packed columns of chelating Sepharose Fast Flow beads (GE Healthcare) charged with nickel sulphate. The columns were washed with at least 5 column volumes of buffer containing 4 M urea, 20 mM Tris pH 7.5, 15 mM imidazole, and proteins were eluted from the Ni-NTA resin with buffer containing 4 M urea, 20 mM Tris pH 7.5, 500 mM imidazole. TEV cleavage of the 6xHis-tag was performed while dialyzing against a buffer containing 2 M urea, 20 mM Tris pH 7.5, 50 mM NaCl, 0.5 mM EDTA, 1 mM DTT overnight at room temperature. After cleavage, the protein solutions were loaded onto Ni-NTA columns to separate the protein of interest from the cleaved His-tag. The fractions containing the flow-through and buffer wash were collected and concentrated using a 3,000 MWCO centrifugal filter (Amicon, Millipore Sigma). For the final purification step, the samples were passed over a S75 Superdex size exclusion column (Cytiva Life Sciences) in 2 M guanidinium chloride, 20 mM MES pH 5.5. The identity of each protein was confirmed via intact mass spectrometry. All proteins were stored in 4 M guanidinium chloride, 20 mM MES pH 5.5 at 4°C. For the −4D+4V and −4D+4N variants, which are highly aggregation prone, storage was performed in 6 M guanidinium chloride. The allW variants, which showed a tendency to phase-separate even during TEV cleavage, were diluted to a minimum volume of 80–100 mL per 1 L of culture and cleaved with TEV protease in 3 M urea, 20 mM Tris pH 7.5, 25 mM NaCl, 0.5 mM EDTA, 1 mM DTT. For the allW variants, size-exclusion chromatography was performed in 4 M guanidinium chloride, 20 mM MES, pH 5.5 to keep the proteins soluble.

### Buffer exchange into native buffer

All variants used in this study showed a tendency to aggregate upon slow buffer exchange through dialysis. Therefore, rapid buffer exchange was performed through use of Zeba spin desalting columns (7K MWCO, ThermoFisher Scientific) or HiTrap desalting columns with Sephadex G-25 resin (Cytiva Life Sciences).

### SDS-PAGE

Denaturing polyacrylamide gel electrophoresis was performed using NuPAGE 4-12% Bis-Tris gradient gels (Invitrogen) and 1X NuPAGE MES SDS running buffer (Invitrogen). Gel staining was performed with SimplyBlue SafeStain (Thermo Fisher Scientific) followed by destaining in water. PageRuler Plus Prestained protein ladder (Thermo Fisher Scientific) was used as a molecular mass ladder for reference.

### Measurement of saturation concentrations

Dilute phase concentrations were determined as a function of temperature (4°C to 20°C) using the sedimentation protocol described previously^71^. Briefly, phase separation was induced by adding NaCl to a final concentration of 150 mM, samples were incubated on temperature blocks for 5-10 minutes, and the dilute and dense phases were separated via centrifugation under pre-equilibrated temperature (21,100 g for 5 min). The protein concentration of the dilute phase was determined by measurements of the absorbance at 280 nm on a UV-Vis spectrophotometer (NanoDrop, ThermoFisher Scientific). For samples with low absorbance values, measurements were performed in a 10 mm pathlength quartz cuvette. All measurements were performed at least in triplicate.

### Differential interference contrast (DIC) microscopy

DIC microscopy was performed on a Nikon Eclipse Widefield microscope or a 3i Marianas spinning disk confocal microscope with a 20 X objective. Samples were prepared by adding NaCl to a final concentration of 150 mM. Protein concentrations were selected such that they were at approximately equal values of supersaturation for the different variants. Images were acquired at room temperature. Protein solution (1.5–3 µL) was sandwiched between a slide and coverslip (1.5 mm thickness) held together by 3M 300 LSE high-temperature double-sided tape (0.34 mm) with holes punched in to accommodate the samples.

### Protein solubility measurements

Samples containing two input protein concentrations in the sub-saturated and super-saturated regimes were prepared in triplicates for each variant. A final NaCl concentration of 150 mM was used, and 0.03% sodium azide (NaN_3_) was added to avoid bacterial contamination during the prolonged incubation. The samples were incubated at 20°C and 600 rpm on a bench-top incubator shaker for 14 days. After the incubation period, the samples were transferred to 1.5 mL ultracentrifuge tubes (Beckman Coulter) and ultracentrifugation was performed at 218,000 g for 40 minutes on a Beckman Coulter Ultracentrifuge (Optima). The soluble protein concentration of the supernatant was measured from the absorbance at 280 nm in a 10 mm pathlength cuvette in a UV-Vis spectrophotometer.

### Confocal Microscopy for ThT localization

To study spatial localization of ThT fluorescence in condensates as a function of time, a Zeiss LSM 780 confocal microscope was used. The samples were prepared by adding ThT (to a final concentration of 20 μM) to the protein samples and inducing phase separation by adding NaCl to a final concentration of 150 mM. The samples (1.5–3 μL) were either sandwiched as described above or placed (15 μL) into uncoated, chambered μ-Slide 15-Well 3D (Ibidi) coverslips, with the wells surrounding the sample wells filled with water to slow down sample evaporation. The plates were sealed with DIC compatible lids. Imaging was performed at different time points using the following settings: 20X objective or 40X water objective; laser excitation at 458 nm, emission at 472–552 nm. Image processing was performed using Fiji.

### Confocal microscopy for imaging condensates

N-terminal labeling was performed in denaturing conditions (4 M guanidinium chloride, 100 mM phosphate buffer, pH 7.0) for the A1-LCD variants using either Oregon Green 488 or Alexa Fluor 647 NHS esters. To generate samples for imaging, labelled protein was spiked into the corresponding unlabeled variant at a molar ratio of either 1:100 or 1:300, for Oregon Green 488 or Alexa Fluor 647, respectively. The protein samples were buffer exchanged into native buffer, followed by induction of phase separation by adding NaCl to a final concentration of 150 mM. For long-term imaging, sample was prepared as follows: a 4 μL drop of sample was sandwiched between a slide and a cover glass separated by two narrow strips of double-sided adhesive tape (3M 300LSE), and the region surrounding the drop was filled with mineral oil to prevent evaporation. The images were collected on a spinning disc confocal microscope (3i Marianas) using laser excitation of 488 nm and 633 nm for Oregon Green 488 and Alexa Fluor 647, respectively.

### Thioflavin T fluorescence assays

Samples of equal volume generated via a 0.8x serial dilution of A1-LCD variants were prepared in the wells of a 96-well clear flat-bottom half-area microplate (Corning or Greiner Bio-One). Just before starting the measurements, phase-separation was induced by adding ThT to a final concentration of 20 μM and NaCl to a final concentration of 150 mM to all wells. Plates were sealed with optically transparent film to prevent evaporation, and data collection was performed at room temperature with 15 seconds of shaking at 500 rpm between readings. ThT fluorescence (excitation 450 nm, emission 510 nm) was monitored every few minutes (1-5 minutes), for 24-36 hours using either the top-reading (black plates) or the bottom-reading mode (clear plates) in a plate reader (CLARIOstar, BMG Labtech). Data analysis and plotting were performed using custom scripts written in R.

### Preparation of fibril seeds

Samples of fibril seeds were prepared as follows: Samples of monomeric D262V A1-LCD of a known concentration and volume were incubated in 40 mM HEPES, pH 7, 150 mM NaCl for 24 hours at 600 rpm until fibril formation. Fibrils were separated by ultracentrifugation at 218,000 g for 40 minutes in a Beckman Coulter Ultracentrifuge (Optima). The concentration of the supernatant was determined, and a specified volume of supernatant was removed. Fibril concentration was determined in terms of monomer equivalents compared to the initial monomer concentration and the leftover concentration in the supernatant. The fibril pellet was resuspended in buffer, vortexed and sonicated to prepare seeds. For the final assay, seeds were used as 10% w/w of the monomeric protein concentration.

### Transmission Electron Microscopy (TEM)

Samples were adsorbed to freshly glow-discharged 300 mesh carbon/formvar-coated grids for electron microscopy by floating support on a drop of sample solution for 10 minutes. Following adsorption, samples were washed three times in ddH2O by passing the grid across a large droplet of water. Grids were wicked almost dry with filter paper then contrasted by floating on a drop of 2% uranyl acetate for 1 minute. Excess stain was wicked off the grid, and grids were allowed to air dry. Samples were imaged in a ThermoFisher Scientific F20 transmission electron microscope operating at 80 kV, and images were recorded on an AMT camera system.

### Prediction of saturation concentrations from the mean-field model

The measured *c*_sc_ values of variants at 4°C were converted into the rescaled *c*_sc_ values referred to as *c*_sc_′ by using the equation described by Bremer et al. ^31^. The predicted values of *c*_sc_′ for these mutants were obtained from the mean-field model derived from the V-shaped plot of rescaled saturation concentrations of A1-LCD variants vs NCPR ^31^. To determine the agreement between them, the measured *c*_sc_′ values were plotted against the predicted *c*_sc_′ values.

### Chemical kinetics modeling

Modeling was based on the following assumptions: (1) The system comprises three species that mimic the dilute, dense (condensate), and fibrillar phases; (2) fibril formation can only occur in the dilute phase, (3) protein exchange occurs between the dilute and dense phases with rate constants controlling the influx (*k*_dil→den_) and efflux (*k*_den→dil_), (4) fibril formation is controlled by a combination of primary nucleation (*k*_1_), secondary nucleation (*k*_2_), and elongation (*k*_long_), and (5) monomer loss (i.e., reversal of elongation, *k*_loss_) is set to zero. Note that the term “phase” is used loosely here because there is no phase boundary, with a delineating interface. Furthermore, we use a chemical kinetics approximation, which means that neither the transport properties within distinct phases nor the relative volumes or volume fractions are constrained in any way. The predictions are interpreted within these limitations of the model. For example, if we assume two coexisting phases, α and β, then the fraction of molecules in each of the phases, denoted as f(α) and f(β) would be written as: f(x)=(c(x)β(x))/c^tot^, where x is α or β, c(x) is the equilibrium concentration in phase x, β(x) is the fraction of the system volume that is occupied by phase x, and c^tot^ is the input concentration. The expressions for f(x) become more complicated when we have three coexisting phases α, β, and γ, which correspond to dilute, dense, and fibrillar phases, respectively. Importantly, we do not have information regarding the steady-state values or time dependencies of each of the β(x) values. Instead, we have information regarding the growth of fibrillar species in the form of the gain in ThT fluorescence as a function of time. Therefore, the model can be used to fit the kinetic traces and make phenomenological predictions, which are then tested using a different set of experiments. The goodness of fit is a necessary albeit insufficient test of the exactness or accuracy of the model. However, the goodness of fit can be used to make predictions, which are testable in separate experiments. These predictions and the tests are described in the main text. In the model, coupled differential equations are used to model the temporal evolution of different species, for different total protein concentrations. These equations, which are written in terms of the concentrations (activities) of proteins in the dilute phase ([dilute]), dense phase ([dense]), fibrils ([fibrils]), and the number of fibrils (num_fibrils_), are based on the work of Meisl et al. ^51^. They are written as follows:

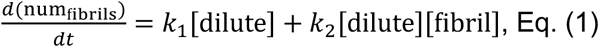

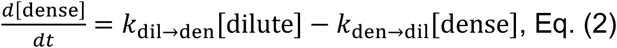

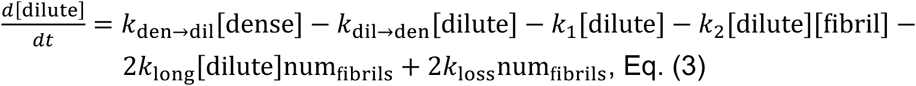

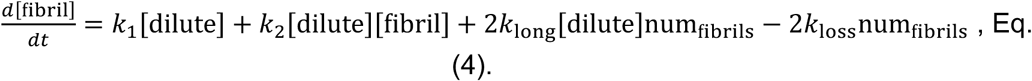

### Fitting experimental ThT traces

Fits were determined for all total concentrations above 17 µM for each of the three replicates corresponding to the sample age of zero days. The parameter fluor_scaling_ was used to convert the ThT fluorescence to protein concentrations using a linear relationship. The known experimental total concentrations were used as the starting concentrations in the model. The parameters *k*_dil→den_ and *k*_den→dil_ were set to zero for concentrations below *c*_sc_ where dense phases (condensates) are absent. Optimization was performed using a least-squares residuals model. Several initial starting parameters were used for the lowest concentration for each construct to minimize the residuals. Then, the best fit parameters from the previous concentration were used as starting points to fit the ThT traces for the next concentration. Fits were performed over the first 500 minutes of the ThT traces unless traces showed a large drop in ThT fluorescence likely due to fibrils accumulating at the bottom of the well and therefore not contributing to the ThT signal. The slopes of the ThT traces were analyzed to extract traces that agreed with this criterion and thus the revised time cutoff for fitting. All initial and extracted parameters, time cutoffs, and fit values for each experimental concentration and replicate are provided.

### Extraction of additional information from chemical kinetics modeling

A time step of 10^-3^ minutes was used. Euler’s method was used to calculate protein concentrations. To determine the effect of titrating *k*_dil→den_ and *k*_den→dil_ on the time to reach 5, 50, and 95 percent fibril, all other parameters were set to be constant. Specifically, *k*_1_ = 1e-8, *k*_2_ = 5e-9, *k*_long_ = 500, and the total protein concentration was set to 200 µM. These values were chosen to be equivalent to extracted parameters from fits of WT and D262N ThT traces at 200 µM. Simulations were performed for a total of 5×10^5^-time steps, and this equates to 500 minutes. To plot protein fractions in the dilute, dense, and fibrillar phases, the extracted best parameters from fits to the ThT traces were utilized. Here, simulations were performed for 5×10^5^-time steps, and this again equates to 500 minutes. Given a total concentration, *c*_total_, the initial dilute phase concentration was set by *c*_total_*k*_den→dil_/(*k*_dil→den_+ *k*_den→dil_) and the initial dense phase concentration was set by *c*_total_*k*_dil→den_/(*k*_dil→den_+ *k*_den→dil_).

### Condensate dissolution assay under flow

Condensates of WT A1-LCD and two variants, allW and allW D262V, prepared in 40 mM HEPES buffer (pH 7.0) and 150 mM NaCl, were flowed into a laminar flow microfluidic chamber (u-Flux) attached to a Lumicks C-trap correlative laser tweezer and confocal fluorescence microscopy system for recording condensate dissolution kinetics. Using Python-based bluelake software (Lumicks), a pressure of ∼1 bar was set to enable the laminar flow of condensates into the microfluidic cell already equilibrated with the sample buffer. Condensates were optically trapped using a laser tweezer operating at 10% trapping laser (1064 nm) power corresponding to ∼100 μW. Fluorescence images of condensates, visualized with either Alexa488 (WT) or Alexa647 (allW and allW D262V) labeled A1-LCD (50 nM), were acquired continuously (0.3 to 1.1 frames per second) to monitor the decay in condensate size as the trapped condensate undergoes dissolution under laminar flow conditions (pressure set at ∼0.6 bar). For analysis of condensate dissolution data, the respective timelapse video was imported into Fiji (v1.54f) and thresholding was performed to binarize the timelapse image sequence to better distinguish the droplet from the background for reliable quantification of condensate area. Using the ‘analyze particles’ function of Fiji, the area of the shrinking condensates was extracted as a function of time. GraphPad Prism 10 was used for generating the plots of condensate size over time and fitting with a stretched exponential function was performed to estimate the dissolution time constant (τ) and stretching exponent β of individual condensates of each A1-LCD variant. For generating representative videos of condensate dissolution, the raw timelapse videos were upscaled utilizing bilinear interpolation, with the aid of Fiji, for better clarity.

### Stress granule disassembly assay

U2OS cells expressing G3BP1-tdTomato from the G3BP1 endogenous locus^1^ were cultured in DMEM with 4.5 g/L glucose, 10% FBS and 2 mM L-glutamine supplement and no antibiotics. Sequences of all constructs used in cellular experiments are shown in **Table S2**. Cells were seeded in 35 mm glass bottom dishes (Thermo Fisher # 150680) at a density of 400,000 cells per dish. The following day, full-length hnRNPA1 constructs with C-terminal FLAG tags, and expressing eGFP proportionately via an intermediate IRES sequence cloned in the pcDNA4/TO/myc-his A backbone, were transfected into the cells (1 μg DNA per dish) using Viafect transfection reagent (Promega) as per the manufacturer’s protocol. 24 hours after transfection, cell nuclei were stained with Hoechst 33342 (2 μg/mL) in imaging media (Fluorobrite DMEM with 10% FBS and 2 mM L-glutamine) for 10 minutes. After replacing the staining media with imaging media, cells were imaged using an Okogawa CSU-W1 spinning disk confocal attached to a Nikon Ti2 Eclipse microscope with a Photometrics Prime 95B camera and using Nikon Elements software (v5.21.02). Conditions were maintained at 37°C and 5% CO2 using a Bold Line Cage Incubator (Okolab) and an objective heater (Bioptechs), excluding when heat stress was applied. Imaging was performed through a Nikon Plan Apo 60× 1.40 NA oil objective with Immersol 518 F (Zeiss; refractive index 1.518), and Perfect Focus 2.0 (Nikon) was engaged for all captures. Heat stress was achieved by increasing the temperature of the objective heater to 43°C, which showed a reproducible effect of stress granule formation after 10-11 minutes. After 1 hour of heat stress, the objective heater temperature was lowered to 37°C to allow the cells to recover, during which time the stress granules gradually disassemble. Time-lapse imaging was performed for a total of 3 hours (1 hour of heat stress and 2 hours of recovery). For different days of experiment, the sequential order of the variants imaged were cycled to randomize the effect of variable wait time before imaging.

### Analysis of stress granule disassembly

20 regions for each sample were imaged simultaneously, resulting in 20 time-lapse videos per day of experiment and construct. The time-lapse videos from each condition for each experiment day were manually reviewed. All cells that (a) have eGFP expression above background fluorescence, (b) show stress granules at the end of the heat stress, and (c) remain viable over the course of the experiment were included in the test set. For the negative control, cells that (a) do not express eGFP above background fluorescence, (b) show stress granules at the end of the heat stress, and (c) and remain viable over the course of the experiment, were picked, 4 cells per time-lapse video, one from each quadrant. Cells were manually tracked to determine the timepoint at which no stress granules are visible. Time-lapse videos were included as long as a few defocused frames did not interfere with an accurate determination of the timepoint at which all stress granules disassemble. The stress granule disassembly timepoints for each construct for three independent days of experiments were combined and analysed using standard survival analysis methods. Kaplan-Meier survival curves that represent the fraction of cells showing stress granules at each timepoint were generated **(Figure 7C,D)**. The survival curves for different variants were compared using log rank testing, and the p-values were adjusted with the Benjamini-Hochberg correction for multiple comparisons ^72^.

### Analysis of stress granule area fraction

For each of the stress granule disassembly experiments, the 20^th^ frame (corresponding to 17 minutes of heat stress) was segmented for stress granules using the pixel classification model using Ilastik v1.4 (https://www.ilastik.org/). Cell boundaries and nuclear boundaries were segmented using Cellpose v2 (https://www.cellpose.org/) using the “cyto2” and “nuclei” models, respectively. Only cells expressing eGFP above background were considered. The stress granule area fraction per cell was calculated as stress granule area / (total cell area – nuclear area). Comparison between variants was done by one-way ANOVA followed by Tukey’s post-hoc test.

### Immunofluorescence microscopy

U2OS cells grown in 24-well plates with a cover glass at the bottom were grown to 80% confluency before transfecting with the prior-mentioned hnRNPA1-FLAG-IRES-eGFP-myc constructs. 24 hours after transfection, the transfection media was replaced with fresh media and the cells allowed to recover for an hour. The transfected cells were subjected to heat stress for 1 hour using a 43°C incubator followed by recovery for 2 hours at 37°C. At the end of each timepoint, cells were taken out, fixed and permeabilized using 4% paraformaldehyde and 0.5% Triton X-100 for 15 minutes. Antigens were blocked with 3% BSA to reduce non-specific antibody binding, followed by incubation with a cocktail of primary antibodies. Anti-PABP rabbit antibody (Abcam AB21060), anti-FLAG mouse antibody (Sigma-Aldrich F3165) and anti-Myc chicken antibody (Thermo Fisher A21281) were used to stain the stress granule marker PABP, hnRNPA1-FLAG and eGFP-myc, respectively. Secondary antibodies used were anti-rabbit Alexa Fluor 647 (Invitrogen A31573) and anti-mouse Alexa Fluor 555 (Invitrogen A21422) and anti-chicken Alexa Fluor 488 (Invitrogen A11039). DAPI was used to stain the nuclei. Cover glasses were mounted on slides using ProLong Gold mounting media (Invitrogen). Fixed cells were imaged on a Zeiss LSM 780 confocal microscope using a 63x oil immersion objective. Excitation lasers used were 405 nm for DAPI, 488 nm for Alexa Fluor 488, 532 nm for Alexa Fluor 555 and 633 nm for Alexa Fluor 647.

### Western blot

U2OS cells were grown in 24-well plates and grown to 80% confluency before transfecting with the hnRNPA1-FLAG-IRES-eGFP-myc constructs. 24 hours after transfection, cells were lysed directly on the plate using 2x Laemmli SDS sample buffer with 100 mM DTT. The cell lysates were heated to 95°C for 10 minutes followed by centrifugation to remove the insoluble fraction. Equal volumes of the supernatant were separated by SDS-PAGE with a 10% polyacrylamide gel. The gel was blotted on a PVDF membrane using wet transfer method, washed and blocked using a standard protocol, and probed with a cocktail of primary antibodies: anti-hnRNPA1 rabbit polyclonal antibody raised against residues 8-42 (Thermo Fisher PA5-79381), anti-GFP-B2 mouse monoclonal antibody (Santa Cruz Biotechnology sc-9996) and anti-β-actin rabbit monoclonal antibody (Cell Signaling Technology 4970S). All primary antibodies were used at 1:1000 dilutions. The secondary antibodies used were anti-Rabbit IRDye800CW (LICOR Bio 926-32213) and anti-mouse IRDye680RD (LICOR Bio 926-68072), both used at 1:5000 dilution. The membranes were imaged on a LICOR Odyssey Fc imager using excitation wavelengths of 800 nm and 700 nm, respectively.

Video S1. Time-lapse movie of a solution of A1-LCD D262N that is supersaturated with respect to condensate formation transitions to fibrils.

Video S2. Time-lapse movie of a solution of A1-LCD D262V that is supersaturated with respect to condensate formation transitions to fibrils.

Video S3. Time-lapse movie of a solution of WT A1-LCD that is supersaturated with respect to condensate formation transitions to fibrils.

Video S4. Time-lapse movie of the dissolution of a WT A1-LCD condensate under flux in the microfluidic flow chamber.

Video S5. Time-lapse movie of the dissolution of an allW A1-LCD condensate under flux in the microfluidic flow chamber.

Video S6. Time-lapse movie of the dissolution of an allW D262V A1-LCD condensate under flux in the microfluidic flow chamber.

## REFERENCES

1. Yang, P., Mathieu, C., Kolaitis, R.M., Zhang, P., Messing, J., Yurtsever, U., Yang, Z., Wu, J., Li, Y., Pan, Q., et al. (2020). G3BP1 Is a Tunable Switch that Triggers Phase Separation to Assemble Stress Granules. Cell 181, 325–345 e328. 10.1016/j.cell.2020.03.046.

2. Sanders, D.W., Kedersha, N., Lee, D.S.W., Strom, A.R., Drake, V., Riback, J.A., Bracha, D., Eeftens, J.M., Iwanicki, A., Wang, A., et al. (2020). Competing Protein-RNA Interaction Networks Control Multiphase Intracellular Organization. Cell 181, 306–324 e328. 10.1016/j.cell.2020.03.050.

3. Guillen-Boixet, J., Kopach, A., Holehouse, A.S., Wittmann, S., Jahnel, M., Schlussler, R., Kim, K., Trussina, I., Wang, J., Mateju, D., et al. (2020). RNA-Induced Conformational Switching and Clustering of G3BP Drive Stress Granule Assembly by Condensation. Cell 181, 346–361 e317. 10.1016/j.cell.2020.03.049.

4. Van Treeck, B., Protter, D.S.W., Matheny, T., Khong, A., Link, C.D., and Parker, R. (2018). RNA self-assembly contributes to stress granule formation and defining the stress granule transcriptome. Proceedings of the National Academy of Sciences 115, 2734–2739. 10.1073/pnas.1800038115.

5. Wang, J., Choi, J.-M., Holehouse, A.S., Lee, H.O., Zhang, X., Jahnel, M., Maharana, S., Lemaitre, R., Pozniakovsky, A., Drechsel, D., et al. (2018). A Molecular Grammar Governing the Driving Forces for Phase Separation of Prion-like RNA Binding Proteins. Cell 174, 688–699.e616. 10.1016/j.cell.2018.06.006.

6. Taylor, J.P. (2015). Multisystem proteinopathy. Intersecting genetics in muscle, bone, and brain degeneration 85, 658–660. 10.1212/wnl.0000000000001862.

7. Ciryam, P., Lambert-Smith, I.A., Bean, D.M., Freer, R., Cid, F., Tartaglia, G.G., Saunders, D.N., Wilson, M.R., Oliver, S.G., Morimoto, R.I., et al. (2017). Spinal motor neuron protein supersaturation patterns are associated with inclusion body formation in ALS. Proceedings of the National Academy of Sciences 114, E3935–E3943. doi:10.1073/pnas.1613854114.

8. Farrawell, N.E., Lambert-Smith, I.A., Warraich, S.T., Blair, I.P., Saunders, D.N., Hatters, D.M., and Yerbury, J.J. (2015). Distinct partitioning of ALS associated TDP-43, FUS and SOD1 mutants into cellular inclusions. Scientific Reports 5, 13416. 10.1038/srep13416.

9. Neumann, M., Bentmann, E., Dormann, D., Jawaid, A., DeJesus-Hernandez, M., Ansorge, O., Roeber, S., Kretzschmar, H.A., Munoz, D.G., Kusaka, H., et al. (2011). FET proteins TAF15 and EWS are selective markers that distinguish FTLD with FUS pathology from amyotrophic lateral sclerosis with FUS mutations. Brain 134, 2595–2609. 10.1093/brain/awr201.

10. Kabashi, E., Valdmanis, P.N., Dion, P., Spiegelman, D., McConkey, B.J., Vande Velde, C., Bouchard, J.P., Lacomblez, L., Pochigaeva, K., Salachas, F., et al. (2008). TARDBP mutations in individuals with sporadic and familial amyotrophic lateral sclerosis. Nature Genetics 40, 572–574. 10.1038/ng.132.

11. Wolozin, B., and Ivanov, P. (2019). Stress granules and neurodegeneration. Nat Rev Neurosci 20, 649–666. 10.1038/s41583-019-0222-5.

12. Bentmann, E., Haass, C., and Dormann, D. (2013). Stress granules in neurodegeneration--lessons learnt from TAR DNA binding protein of 43 kDa and fused in sarcoma. FEBS J 280, 4348–4370. 10.1111/febs.12287.

13. Kim, H.J., Kim, N.C., Wang, Y.D., Scarborough, E.A., Moore, J., Diaz, Z., MacLea, K.S., Freibaum, B., Li, S., Molliex, A., et al. (2013). Mutations in prion-like domains in hnRNPA2B1 and hnRNPA1 cause multisystem proteinopathy and ALS. Nature 495, 467–473. 10.1038/nature11922.

14. Mackenzie, I.R., Nicholson, A.M., Sarkar, M., Messing, J., Purice, M.D., Pottier, C., Annu, K., Baker, M., Perkerson, R.B., Kurti, A., et al. (2017). TIA1 Mutations in Amyotrophic Lateral Sclerosis and Frontotemporal Dementia Promote Phase Separation and Alter Stress Granule Dynamics. Neuron 95, 808–816 e809. 10.1016/j.neuron.2017.07.025.

15. Patel, A., Lee, H.O., Jawerth, L., Maharana, S., Jahnel, M., Hein, M.Y., Stoynov, S., Mahamid, J., Saha, S., Franzmann, T.M., et al. (2015). A Liquid-to-Solid Phase Transition of the ALS Protein FUS Accelerated by Disease Mutation. Cell 162, 1066–1077. 10.1016/j.cell.2015.07.047.

16. Nomura, T., Watanabe, S., Kaneko, K., Yamanaka, K., Nukina, N., and Furukawa, Y. (2014). Intranuclear aggregation of mutant FUS/TLS as a molecular pathomechanism of amyotrophic lateral sclerosis. Journal of Biological Chemistry 289, 1192–1202. 10.1074/jbc.M113.516492.

17. Li, Y.R., King, O.D., Shorter, J., and Gitler, A.D. (2013). Stress granules as crucibles of ALS pathogenesis. Journal of Cell Biology 201, 361–372. 10.1083/jcb.201302044.

18. Ranganathan, S., and Shakhnovich, E. (2022). The physics of liquid-to-solid transitions in multi-domain protein condensates. Biophysical Journal 121, 2751–2766. 10.1016/j.bpj.2022.06.013.

19. Lu, S., Hu, J., Arogundade, O.A., Goginashvili, A., Vazquez-Sanchez, S., Diedrich, J.K., Gu, J., Blum, J., Oung, S., Ye, Q., et al. (2022). Heat-shock chaperone HSPB1 regulates cytoplasmic TDP-43 phase separation and liquid-to-gel transition. Nature Cell Biology 24, 1378–1393. 10.1038/s41556-022-00988-8.

20. Linsenmeier, M., Faltova, L., Morelli, C., Capasso Palmiero, U., Seiffert, C., Kuffner, A.M., Pinotsi, D., Zhou, J., Mezzenga, R., and Arosio, P. (2023). The interface of condensates of the hnRNPA1 low-complexity domain promotes formation of amyloid fibrils. Nature Chemistry 15, 1340–1349. 10.1038/s41557-023-01289-9.

21. Shen, Y., Chen, A., Wang, W., Shen, Y., Ruggeri, F.S., Aime, S., Wang, Z., Qamar, S., Espinosa, J.R., Garaizar, A., et al. (2023). The liquid-to-solid transition of FUS is promoted by the condensate surface. Proceedings of the National Academy of Sciences 120, e2301366120. 10.1073/pnas.2301366120.

22. Choi, C.-H., Lee, D.S.W., Sanders, D.W., and Brangwynne, C.P. (2024). Condensate interfaces can accelerate protein aggregation. Biophysical Journal 123, 1404–1413. 10.1016/j.bpj.2023.10.009.

23. Sear, R.P. (2007). Nucleation: theory and applications to protein solutions and colloidal suspensions. Journal of Physics: Condensed Matter 19, 033101. 10.1088/0953-8984/19/3/033101.

24. Michaels, T.C.T., Qian, D., Šarić, A., Vendruscolo, M., Linse, S., and Knowles, T.P.J. (2023). Amyloid formation as a protein phase transition. Nature Reviews Physics 5, 379–397. 10.1038/s42254-023-00598-9.

25. Crick, S.L., Ruff, K.M., Garai, K., Frieden, C., and Pappu, R.V. (2013). Unmasking the roles of N- and C-terminal flanking sequences from exon 1 of huntingtin as modulators of polyglutamine aggregation. Proceedings of the National Academy of Sciences 110, 20075–20080. 10.1073/pnas.1320626110.

26. Posey, A.E., Ruff, K.M., Harmon, T.S., Crick, S.L., Li, A., Diamond, M.I., and Pappu, R.V. (2018). Profilin reduces aggregation and phase separation of huntingtin N-terminal fragments by preferentially binding to soluble monomers and oligomers. Journal of Biological Chemistry 293, 3734–3746. 10.1074/jbc.RA117.000357.

27. Ruff, K.M., Choi, Y.H., Cox, D., Ormsby, A.R., Myung, Y., Ascher, D.B., Radford, S.E., Pappu, R.V., and Hatters, D.M. (2022). Sequence grammar underlying the unfolding and phase separation of globular proteins. Mol Cell 82, 3193–3208 e3198. 10.1016/j.molcel.2022.06.024.

28. Garai, K., Sahoo, B., Sengupta, P., and Maiti, S. (2008). Quasihomogeneous nucleation of amyloid beta yields numerical bounds for the critical radius, the surface tension, and the free energy barrier for nucleus formation. The Journal of Chemical Physics 128. 10.1063/1.2822322.

29. Molliex, A., Temirov, J., Lee, J., Coughlin, M., Kanagaraj, A.P., Kim, H.J., Mittag, T., and Taylor, J.P. (2015). Phase separation by low complexity domains promotes stress granule assembly and drives pathological fibrillization. Cell 163, 123–133. 10.1016/j.cell.2015.09.015.

30. Martin, E.W., Holehouse, A.S., Peran, I., Farag, M., Incicco, J.J., Bremer, A., Grace, C.R., Soranno, A., Pappu, R.V., and Mittag, T. (2020). Valence and patterning of aromatic residues determine the phase behavior of prion-like domains. Science 367, 694–699. 10.1126/science.aaw8653.

31. Bremer, A., Farag, M., Borcherds, W.M., Peran, I., Martin, E.W., Pappu, R.V., and Mittag, T. (2022). Deciphering how naturally occurring sequence features impact the phase behaviours of disordered prion-like domains. Nature Chemistry 14, 196–207. 10.1038/s41557-021-00840-w.

32. Farag, M., Cohen, S.R., Borcherds, W.M., Bremer, A., Mittag, T., and Pappu, R.V. (2022). Condensates formed by prion-like low-complexity domains have small-world network structures and interfaces defined by expanded conformations. Nature Communications 13, 7722. 10.1038/s41467-022-35370-7.

33. Harmon, T.S., Holehouse, A.S., Rosen, M.K., and Pappu, R.V. (2017). Intrinsically disordered linkers determine the interplay between phase separation and gelation in multivalent proteins. Elife 6. 10.7554/eLife.30294.

34. Buell, A.K., Dobson, C.M., and Knowles, T.P.J. (2014). The physical chemistry of the amyloid phenomenon: thermodynamics and kinetics of filamentous protein aggregation. Essays in Biochemistry 56, 11–39. 10.1042/bse0560011.

35. Eisenberg, D., and Jucker, M. (2012). The amyloid state of proteins in human diseases. Cell 148, 1188–1203. 10.1016/j.cell.2012.02.022.

36. Gui, X., Luo, F., Li, Y., Zhou, H., Qin, Z., Liu, Z., Gu, J., Xie, M., Zhao, K., Dai, B., et al. (2019). Structural basis for reversible amyloids of hnRNPA1 elucidates their role in stress granule assembly. Nature Communications 10, 2006. 10.1038/s41467-019-09902-7.

37. Wilkinson, M., Xu, Y., Thacker, D., Taylor, A.I.P., Fisher, D.G., Gallardo, R.U., Radford, S.E., and Ranson, N.A. (2023). Structural evolution of fibril polymorphs during amyloid assembly. Cell 186, 5798–5811.e5726. 10.1016/j.cell.2023.11.025.

38. Ostwald, W. (1897). Studien uber die Bildung und Umwandlung fester Körper. Zeitschrift für Physikalische Chemie 22, 289–330.

39. Naiki, H., Higuchi, K., Hosokawa, M., and Takeda, T. (1989). Fluorometric determination of amyloid fibrils in vitro using the fluorescent dye, thioflavin T1. Analytical Biochemistry 177, 244–249. 10.1016/0003-2697(89)90046-8.

40. Oxtoby, D.W. (1998). Nucleation of First-Order Phase Transitions. Accounts of Chemical Research 31, 91–97. 10.1021/ar9702278.

41. Ferrone, F. (1999). Analysis of protein aggregation kinetics. Methods Enzymol 309, 256–274. 10.1016/s0076-6879(99)09019-9.

42. Xue, W.F., Homans, S.W., and Radford, S.E. (2008). Systematic analysis of nucleation-dependent polymerization reveals new insights into the mechanism of amyloid self-assembly. Proceedings of the National Academy of Sciences 105, 8926–8931. 10.1073/pnas.0711664105.

43. Arosio, P., Knowles, T.P., and Linse, S. (2015). On the lag phase in amyloid fibril formation. Phys Chem Chem Phys 17, 7606–7618. 10.1039/c4cp05563b.

44. Lipiński, W.P., Visser, B.S., Robu, I., Fakhree, M.A.A., Lindhoud, S., Claessens, M.M.A.E., and Spruijt, E. (2022). Biomolecular condensates can both accelerate and suppress aggregation of α-synuclein. Science Advances 8, eabq6495. 10.1126/sciadv.abq6495.

45. Emmanouilidis, L., Bartalucci, E., Kan, Y., Ijavi, M., Pérez, E., Afanasyev, P., Boehringer, D., Zehnder, J., Parekh, S.H., Bonn, M., et al. (2023). A solid beta-sheet structure is formed at the surface of FUS liquid droplets during aging. Nature Chemical Biology 20, 1044–1052. 10.1038/s41589-024-01573-w.

46. He, C., Wu, C.Y., Li, W., and Xu, K. (2023). Multidimensional super-resolution microscopy unveils nanoscale surface aggregates in the aging of FUS condensates. Journal of the American Chemical Society 145, 24240–24248. 10.1101/2023.07.12.548239.

47. Pantoja-Uceda, D., Stuani, C., Laurents, D.V., McDermott, A.E., Buratti, E., and Mompean, M. (2021). Phe-Gly motifs drive fibrillization of TDP-43’s prion-like domain condensates. PLoS Biology 19, e3001198. 10.1371/journal.pbio.3001198.

48. Maltseva, D., Chatterjee, S., Yu, C.C., Brzezinski, M., Nagata, Y., Gonella, G., Murthy, A.C., Stachowiak, J.C., Fawzi, N.L., Parekh, S.H., and Bonn, M. (2023). Fibril formation and ordering of disordered FUS LC driven by hydrophobic interactions. Nature Chemistry 15, 1146–1154. 10.1038/s41557-023-01221-1.

49. Mukherjee, P., and Ganai, S. (2023). Thioflavin-T: A Quantum Yield-Based Molecular Viscometer for Glycerol-Monohydroxy Alcohol Mixtures. ACS Omega 8, 36604–36613. 10.1021/acsomega.3c06428.

50. Knowles, T.P.J., Waudby, C.A., Devlin, G.L., Cohen, S.I.A., Aguzzi, A., Vendruscolo, M., Terentjev, E.M., Welland, M.E., and Dobson, C.M. (2009). An Analytical Solution to the Kinetics of Breakable Filament Assembly. Science 326, 1533–1537. 10.1126/science.1178250.

51. Meisl, G., Kirkegaard, J.B., Arosio, P., Michaels, T.C., Vendruscolo, M., Dobson, C.M., Linse, S., and Knowles, T.P. (2016). Molecular mechanisms of protein aggregation from global fitting of kinetic models. Nature Protocols 11, 252–272. 10.1038/nprot.2016.010.

52. Michaels, T.C.T., Šarić, A., Habchi, J., Chia, S., Meisl, G., Vendruscolo, M., Dobson, C.M., and Knowles, T.P.J. (2018). Chemical Kinetics for Bridging Molecular Mechanisms and Macroscopic Measurements of Amyloid Fibril Formation. Annual Review of Physical Chemistry 69, 273–298. 10.1146/annurev-physchem-050317-021322.

53. Buell, A.K., Gaspar, R., Kaminski, C.F., Knowles, T.P.J., Linse, S., Meisl, G., Sparr, E., and Young, L. (2017). Secondary nucleation of monomers on fibril surface dominates α-synuclein aggregation and provides autocatalytic amyloid amplification. Quarterly Reviews of Biophysics 50, e6, e6. 10.1017/S0033583516000172.

54. Alshareedah, I., Borcherds, W.M., Cohen, S.R., Singh, A., Posey, A.E., Farag, M., Bremer, A., Strout, G.W., Tomares, D.T., Pappu, R.V., et al. (2024). Sequence-specific interactions determine viscoelasticity and ageing dynamics of protein condensates. Nature Physics 20, 1482–1491. 10.1038/s41567-024-02558-1.

55. Kar, M., Dar, F., Welsh, T.J., Vogel, L.T., Kühnemuth, R., Majumdar, A., Krainer, G., Franzmann, T.M., Alberti, S., Seidel, C.A.M., et al. (2022). Phase-separating RNA-binding proteins form heterogeneous distributions of clusters in subsaturated solutions. Proceedings of the National Academy of Sciences 119, e2202222119. doi:10.1073/pnas.2202222119.

56. Wu, T., King, M.R., Qiu, Y., Farag, M., Pappu, R.V., and Lew, M.D. (2024). Single fluorogen imaging reveals distinct environmental and structural features of biomolecular condensates. Nature Physics In press. 10.1038/s41567-025-02827-7.

57. von Bülow, S., Tesei, G., and Lindorff-Larsen, K. (2024). Prediction of phase separation propensities of disordered proteins from sequence. bioRxiv, 2024.2006.2003.597109. 10.1101/2024.06.03.597109.

58. Erkamp, N.A., Farag, M., Qiu, Y., Qian, D., Sneideris, T., Wu, T., Welsh, T.J., Ausserwӧger, H., Krug, T.J., Chauhan, G., et al. (2024). Differential interactions determine anisotropies at interfaces of RNA-based biomolecular condensates. Nature Communications In press. 10.1101/2024.08.19.608662.

59. Cohen, S.R., Banerjee, P.R., and Pappu, R.V. (2024). Direct computations of viscoelastic moduli of biomolecular condensates. The Journal of Chemical Physics 161. 10.1063/5.0223001.

60. Kawasaki, Y., Watanabe, H., and Uneyama, T. (2011). A Note for Kohlrausch-Williams-Watts Relaxation Function. Nihon Reoroji Gakkaishi 39, 127–131. 10.1678/rheology.39.127.

61. Williams, G., and Watts, D.C. (1970). Non-symmetrical dielectric relaxation behaviour arising from a simple empirical decay function. Transactions of the Faraday Society 66, 80–85. 10.1039/TF9706600080.

62. Pappu, R.V., Cohen, S.R., Dar, F., Farag, M., and Kar, M. (2023). Phase Transitions of Associative Biomacromolecules. Chemical Reviews 123, 8945–8987. 10.1021/acs.chemrev.2c00814.

63. Beijer, D., Kim, H.J., Guo, L., O’Donovan, K., Mademan, I., Deconinck, T., Van Schil, K., Fare, C.M., Drake, L.E., Ford, A.F., et al. (2021). Characterization of HNRNPA1 mutations defines diversity in pathogenic mechanisms and clinical presentation. Journal of Clinical Investigation Insight 6. 10.1172/jci.insight.148363.

64. Lee, Y.J., and Rio, D.C. (2024). A mutation in the low-complexity domain of splicing factor hnRNPA1 linked to amyotrophic lateral sclerosis disrupts distinct neuronal RNA splicing networks. Genes Dev 38, 11–30. 10.1101/gad.351104.123.

65. Mathieu, C., Pappu, R.V., and Taylor, J.P. (2020). Beyond aggregation: Pathological phase transitions in neurodegenerative disease. Science 370, 56–60. 10.1126/science.abb8032.

66. Otte, C.G., Fortuna, T.R., Mann, J.R., Gleixner, A.M., Ramesh, N., Pyles, N.J., Pandey, U.B., and Donnelly, C.J. (2020). Optogenetic TDP-43 nucleation induces persistent insoluble species and progressive motor dysfunction in vivo. Neurobiology of Disease 146, 105078. 10.1016/j.nbd.2020.105078.

67. Yan, X., Kuster, D., Mohanty, P., Nijssen, J., Pombo-Garcia, K., Rizuan, A., Franzmann, T.M., Sergeeva, A., Passos, P.M., George, L., et al. (2024). Intra-condensate demixing of TDP-43 inside stress granules generates pathological aggregates. bioRxiv. 10.1101/2024.01.23.576837.

68. Garcia-Cabau, C., Bartomeu, A., Tesei, G., Cheung, K.C., Pose-Utrilla, J., Pico, S., Balaceanu, A., Duran-Arque, B., Fernandez-Alfara, M., Martin, J., et al. (2024). Mis-splicing of a neuronal microexon promotes CPEB4 aggregation in ASD. Nature. 10.1038/s41586-024-08289-w.

69. Folkmann, A.W., Putnam, A., Lee, C.F., and Seydoux, G. (2021). Regulation of biomolecular condensates by interfacial protein clusters. Science 373, 1218–1224. 10.1126/science.abg7071.

70. Studier, F.W. (2005). Protein production by auto-induction in high density shaking cultures. Protein Expr Purif 41, 207–234. 10.1016/j.pep.2005.01.016.

71. Milkovic, N.M., and Mittag, T. (2020). Determination of Protein Phase Diagrams by Centrifugation. In Intrinsically Disordered Proteins: Methods and Protocols, B.B. Kragelund, and K. Skriver, eds. (Springer US), pp. 685–702. 10.1007/978-1-0716-0524-0_35.

72. Benjamini, Y., and Hochberg, Y. (1995). Controlling the False Discovery Rate: A Practical and Powerful Approach to Multiple Testing. Journal of the Royal Statistical Society: Series B 57, 289–300.

