## Supplemental Information for "Tunable metastability of condensates reconciles their dual roles in amyloid fibril formation"

### Metastable condensates suppress conversion to amyloid fibrils

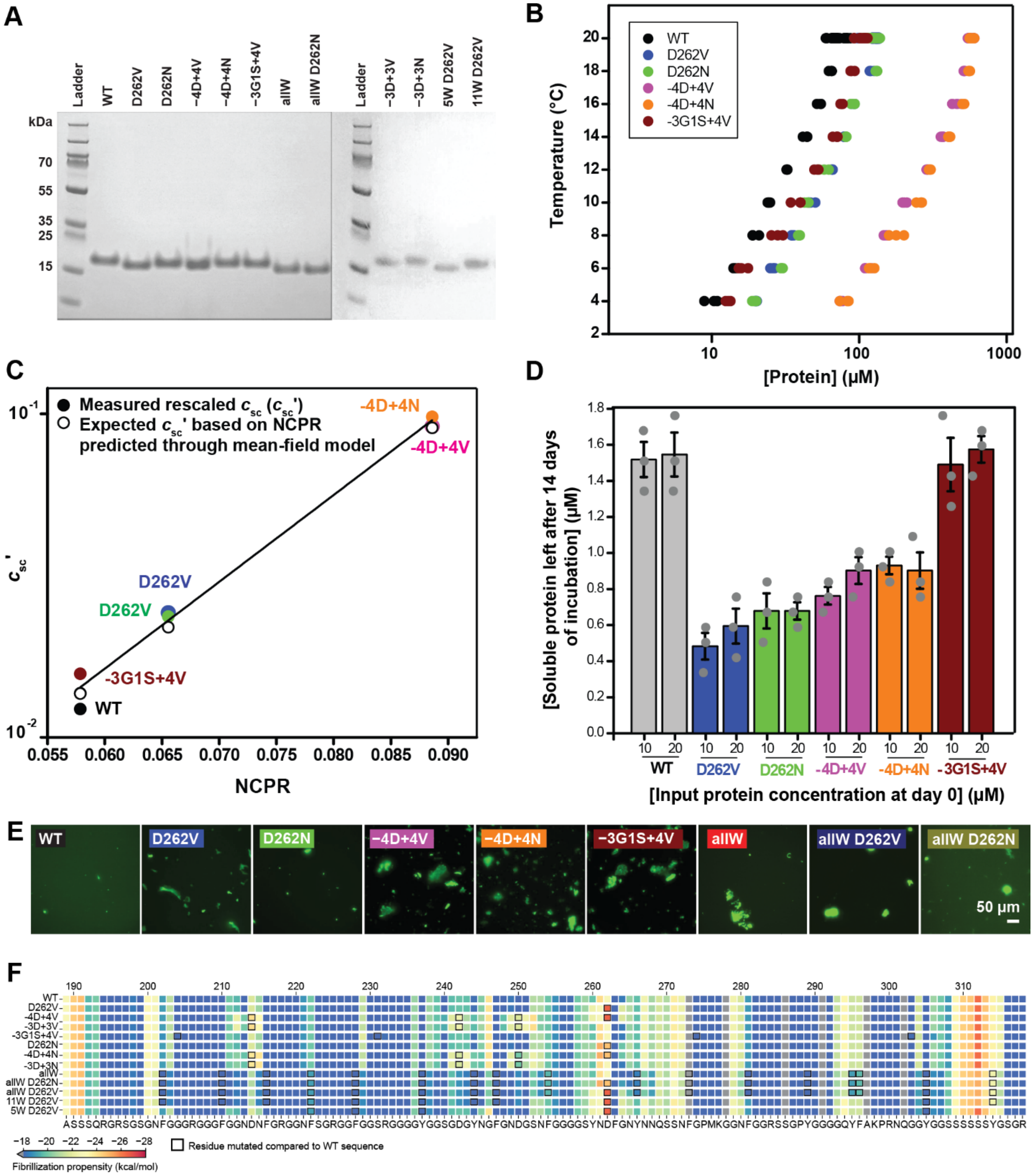


**Figure S1. Pathogenic mutations destabilize condensates, related to Figure 1. (A)** SDS-PAGE analysis of purified A1-LCD variants used in the study. **(B)** Binodals of WT, pathogenic mutants and designed variants as a function of temperature in 40 mM HEPES, 150 mM NaCl, pH 7.0. **(C)** Plot showing a comparison of the rescaled values derived from experimental saturation concentration values for phase-separation ($c_{sc}'$) of the variants used in this study with the predicted $c_{sc}'$ values obtained from the mean-field stickers-and-spacers model developed to describe the phase behavior of the A1-LCD [1]. **(D)** Concentration of soluble protein left in the supernatant after 14 days of incubation, when using different input concentrations below $c_{sc}$. The resulting supernatant concentration are identical within error. Data from individual measurements are shown. **(E)** Fluorescence micrographs depicting ThT-stained pellets obtained after ultracentrifugation of samples used for soluble protein estimation below $c_{sc}$ (see Fig. 1D). **(F)** Predictions from zipperDB[2] showing fibrillization propensity scores as energies in units of kcal/mol for WT A1-LCD and all variants. While Rosetta energies change considerably around position D262 upon introduction of pathogenic mutations, the overall sequence-resolved Rosetta energies change negligibly across the entirety of the sequence in the designed variants. Scores are given for six residue peptides, and the middle ($4^{th}$) residue of the WT A1-LCD peptide sequence is shown across the bottom. Peptides with Rosetta energy scores below −23 kcal/mol (reddish colors) are likely to have high fibrillization propensity. Undefined scores from ZipperDB are set to grey.

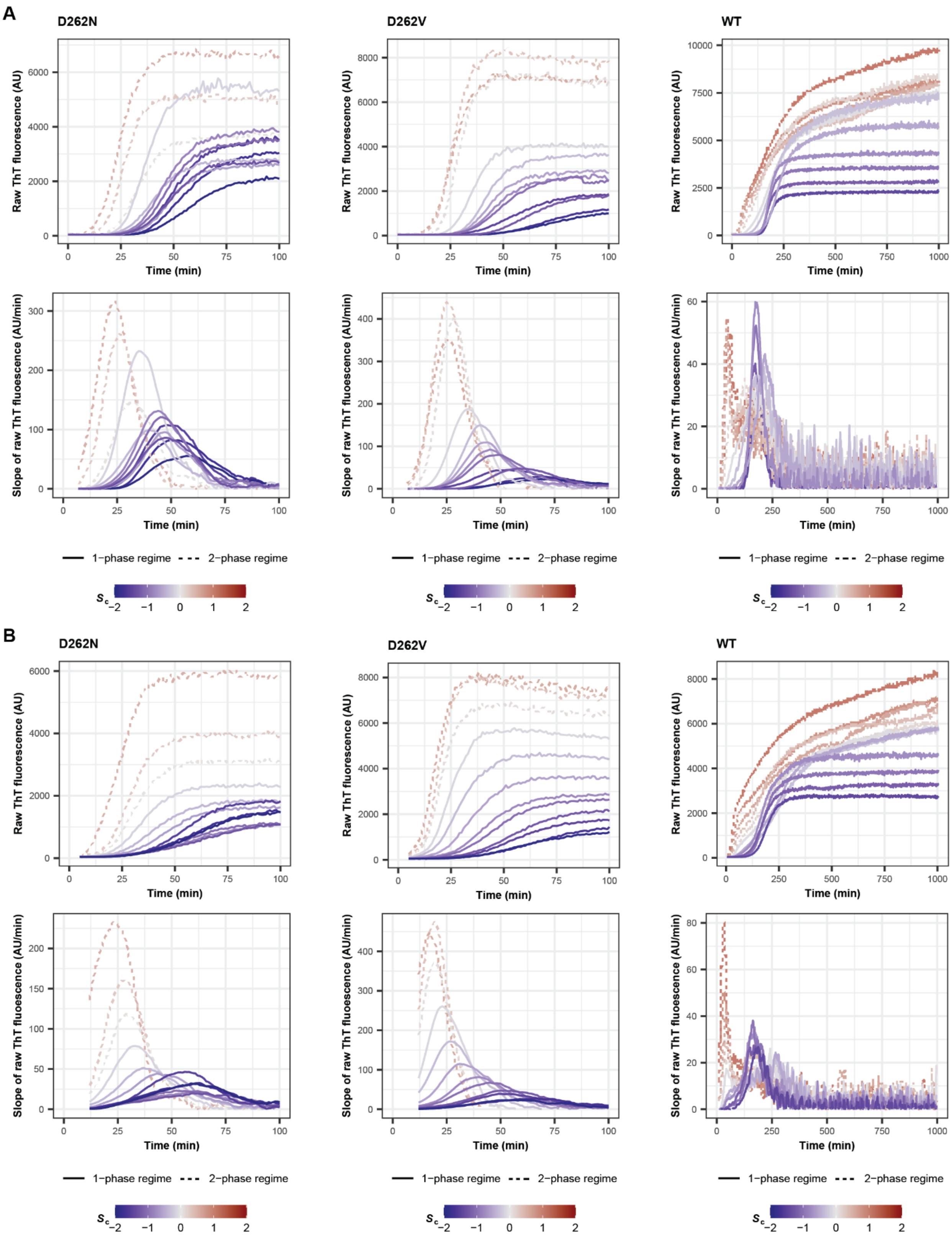
A
D262N
D262V
WT
Raw ThT fluorescence (AU)
Time (min)
Slope of raw ThT fluorescence (AU/min)
1−phase regime
2−phase regime
$S_c$
B
D262N
D262V
WT

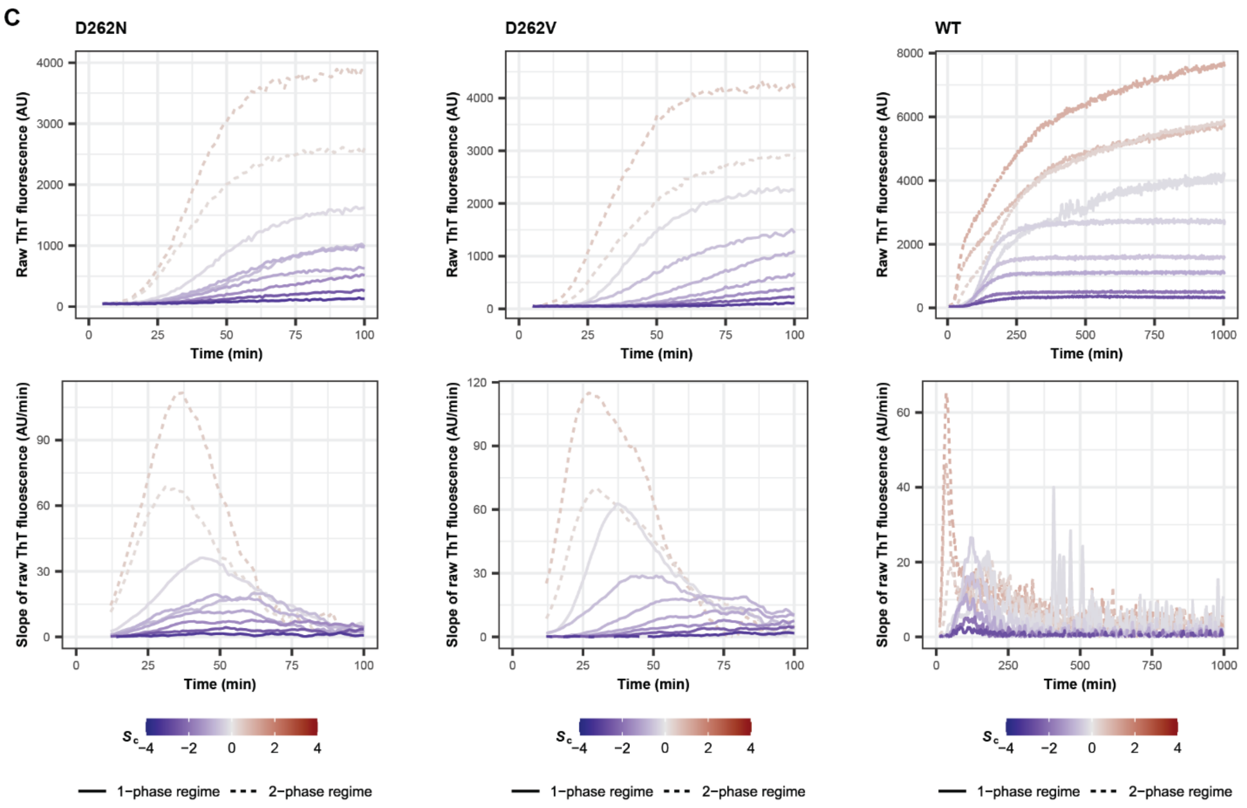
C
D262N
D262V
WT
Raw ThT fluorescence (AU)
Slope of raw ThT fluorescence (AU/min)
Time (min)
Sc
1−phase regime
2−phase regime

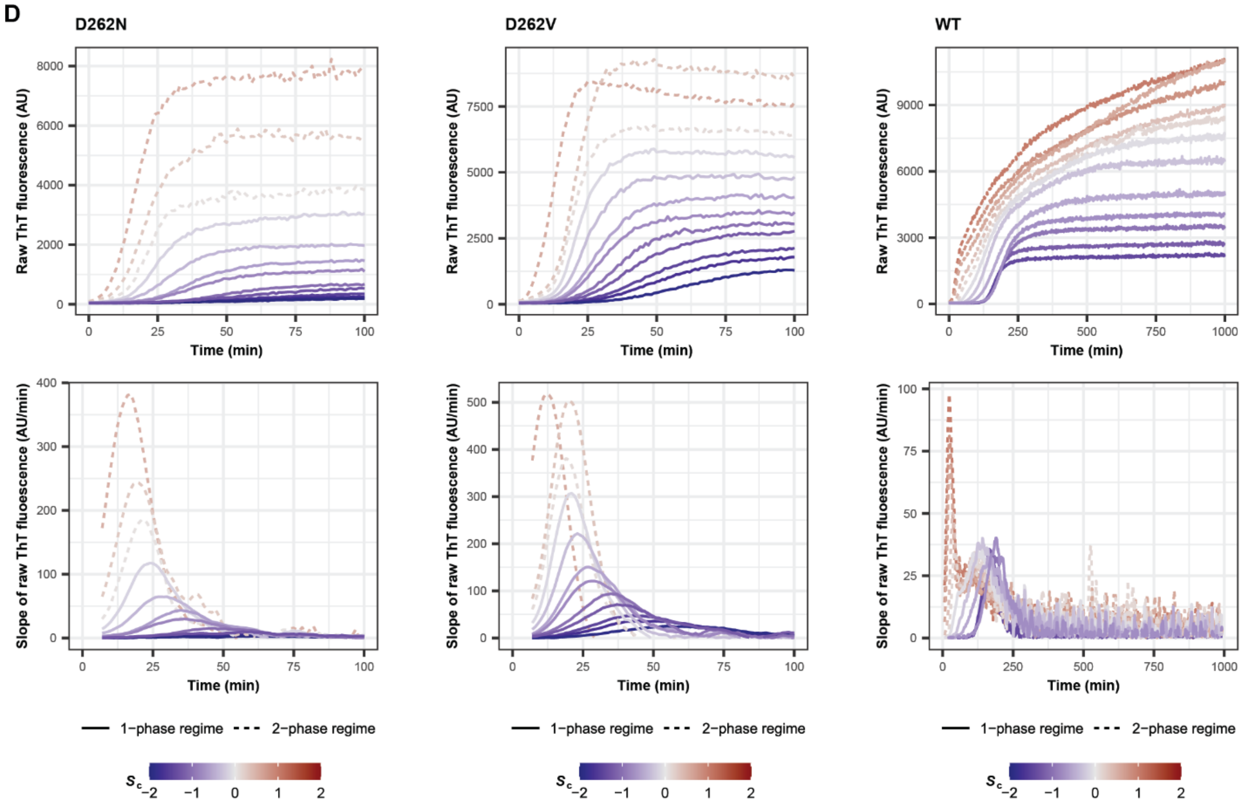
D
D262N
D262V
WT
Raw ThT fluorescence (AU)
Slope of raw ThT fluorescence (AU/min)
Time (min)
1−phase regime
2−phase regime
Sc

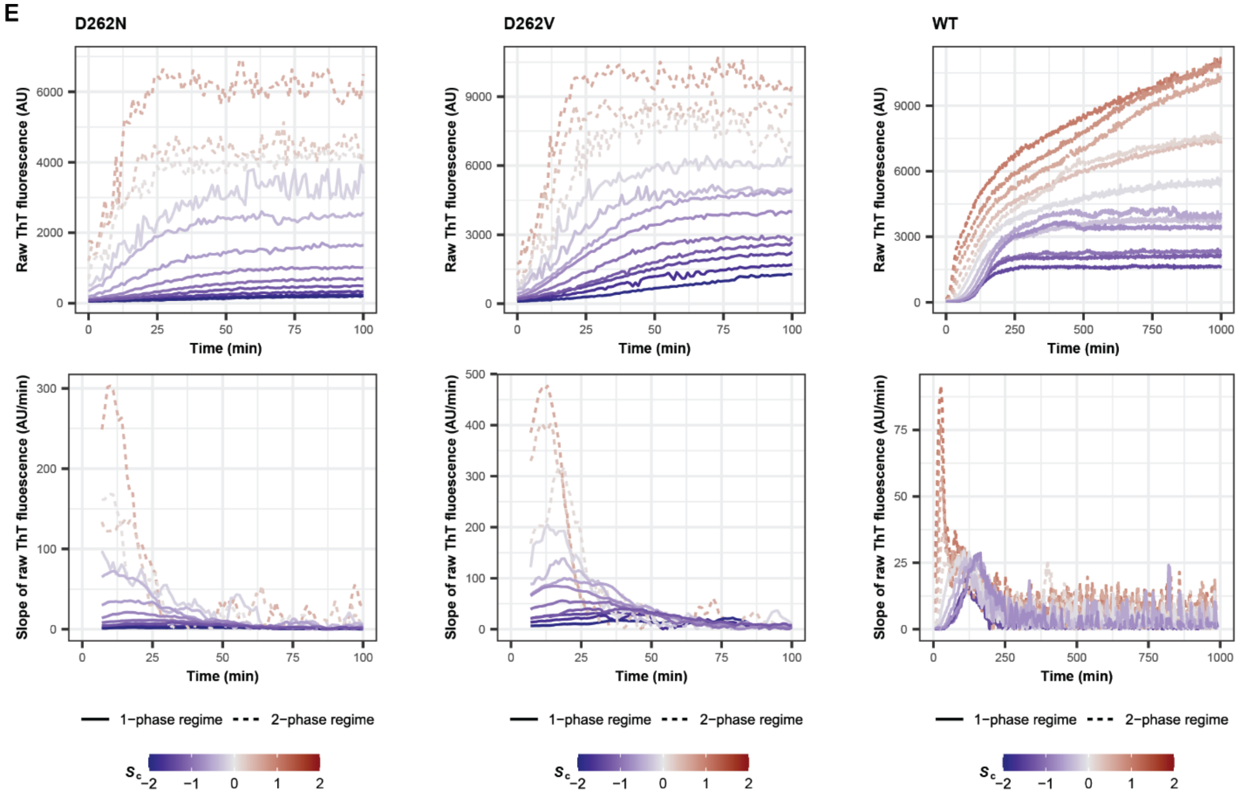


**Figure S2. Kinetics of fibril formation for A1-LCD variants D262N, D262V and WT, related to Figure 2**. **(A-E)** (Top) ThT fluorescence as a function of time across a range of total protein concentrations ($c_{\text{total}}$) that are in the subsaturated (degree of saturation relative to condensates $S_c < 0$, blue solid lines) and supersaturated regimes ($S_c > 0$, red dashed lines) with respect to condensates $\left(S_c = \ln\frac{c_{\text{total}}}{c_{\text{sc}}}\right)$. (Bottom) Rate of increase of ThT fluorescence, i.e., the slope, as a function of time. Data from five independent experiments are shown where data in A–C represent experiments performed on the day of buffer exchange of the protein sample from denaturing conditions into a native buffer, and data in D and E are from experiments performed 1 or 2 days after buffer exchange, respectively. To start the observation time of the experiment, NaCl was added to the protein solutions to 150 mM final concentration.

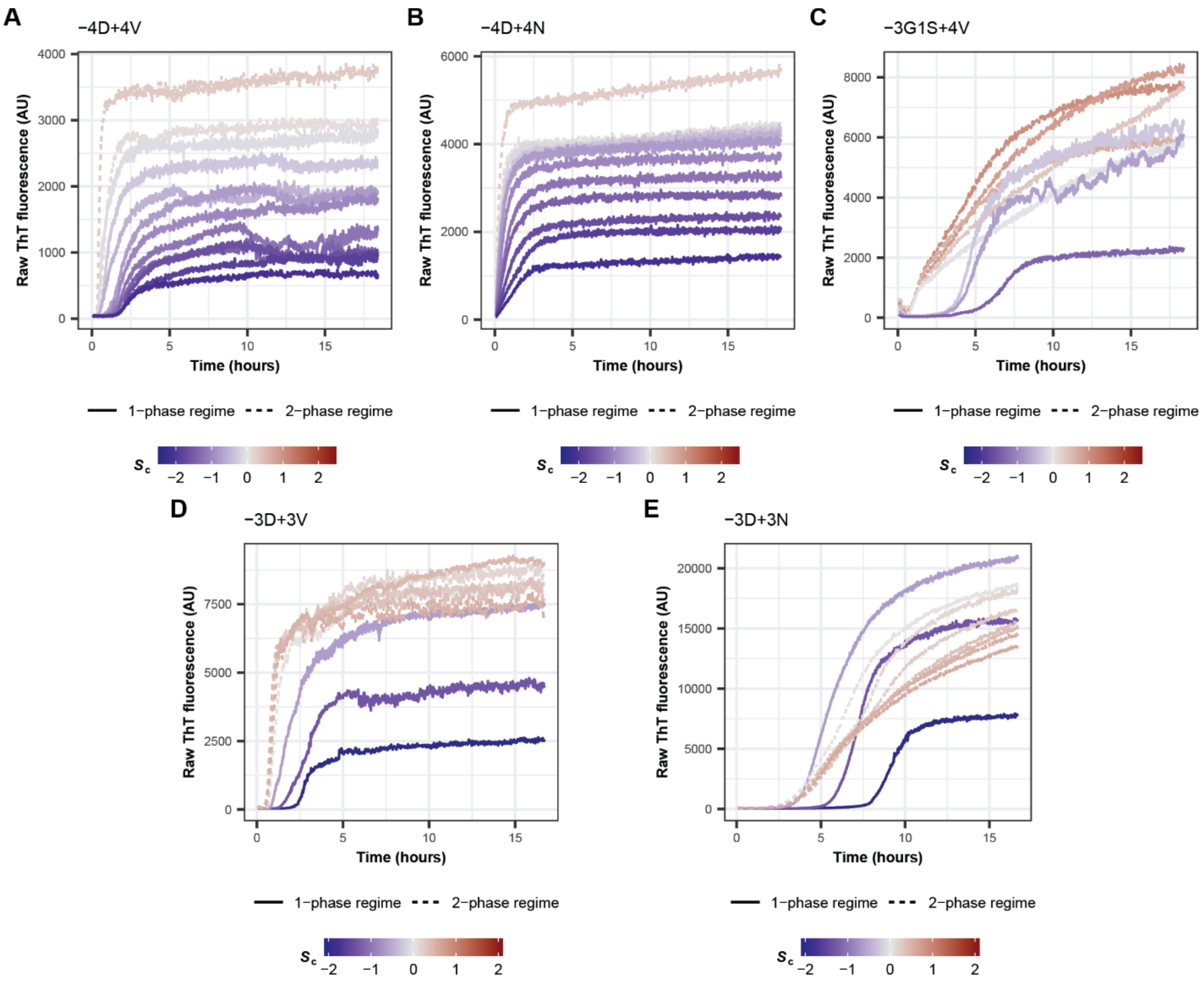


**Figure S3. The effect of designed mutants on kinetics of fibril formation depends on their metastability, related to Figure 2.** Kinetics of fibril formation of **(A)** −4D+4V, **(B)** −4D+4N, **(C)** −3G1S+4V, **(D)** −3D+3V and **(E)** −3D+4N variants monitored by ThT fluorescence. ThT fluorescence as a function of time across a range of total protein concentrations ($c_{\text{total}}$) that are in the subsaturated (degree of saturation relative to condensates $S_c < 0$, blue solid lines) and supersaturated regimes ($S_c > 0$, red dashed lines) with respect to condensates $\left(S_c = \ln\frac{c_{\text{total}}}{c_{\text{sc}}}\right)$.

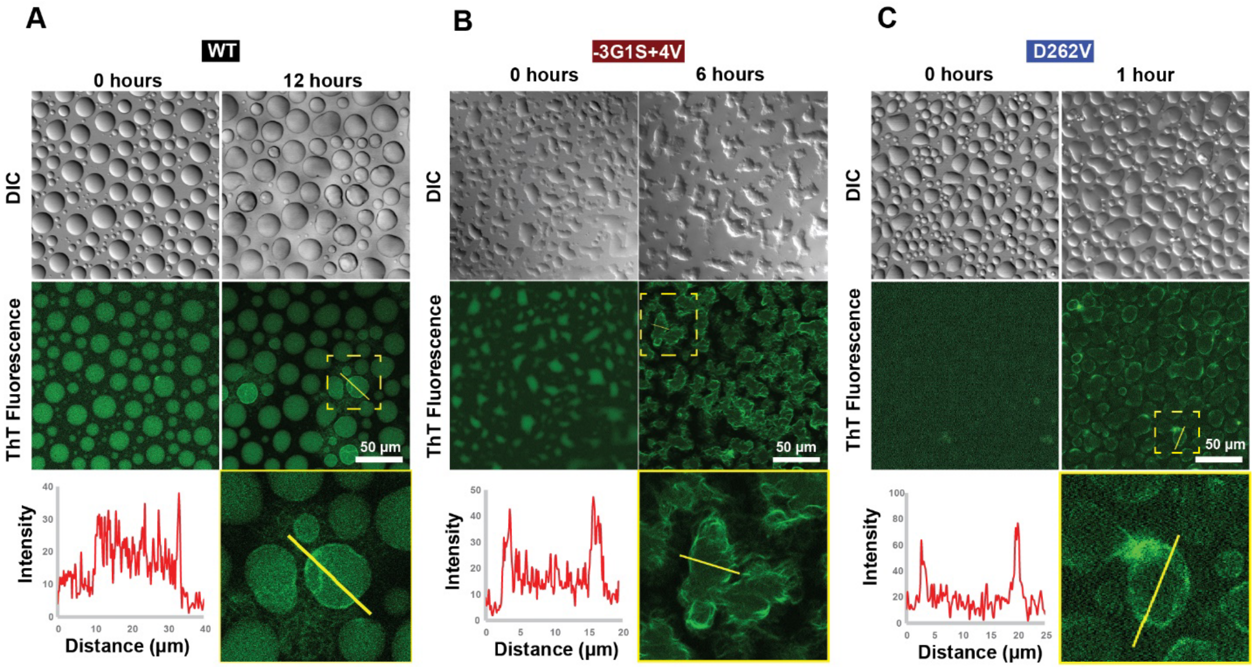


**Figure S4. Condensate interfaces can nucleate fibril formation, related to Figure 2. (A, B, C)** DIC and fluorescence microscopy images to monitor changes in **(A)** WT A1-LCD, **(B)** −3G1S+4V and **(C)** D262V condensate morphology and ThT fluorescence. Inset indicates the portion of fluorescence micrograph shown in the bottom right panel, zoomed in to illustrate the difference in ThT fluorescence intensity between the interface and the interior of the condensates. The line plots show the ThT fluorescence intensity along the yellow line depicted in the image. These data suggest that fibrils nucleate preferentially at interfaces rather than the interiors of condensates.

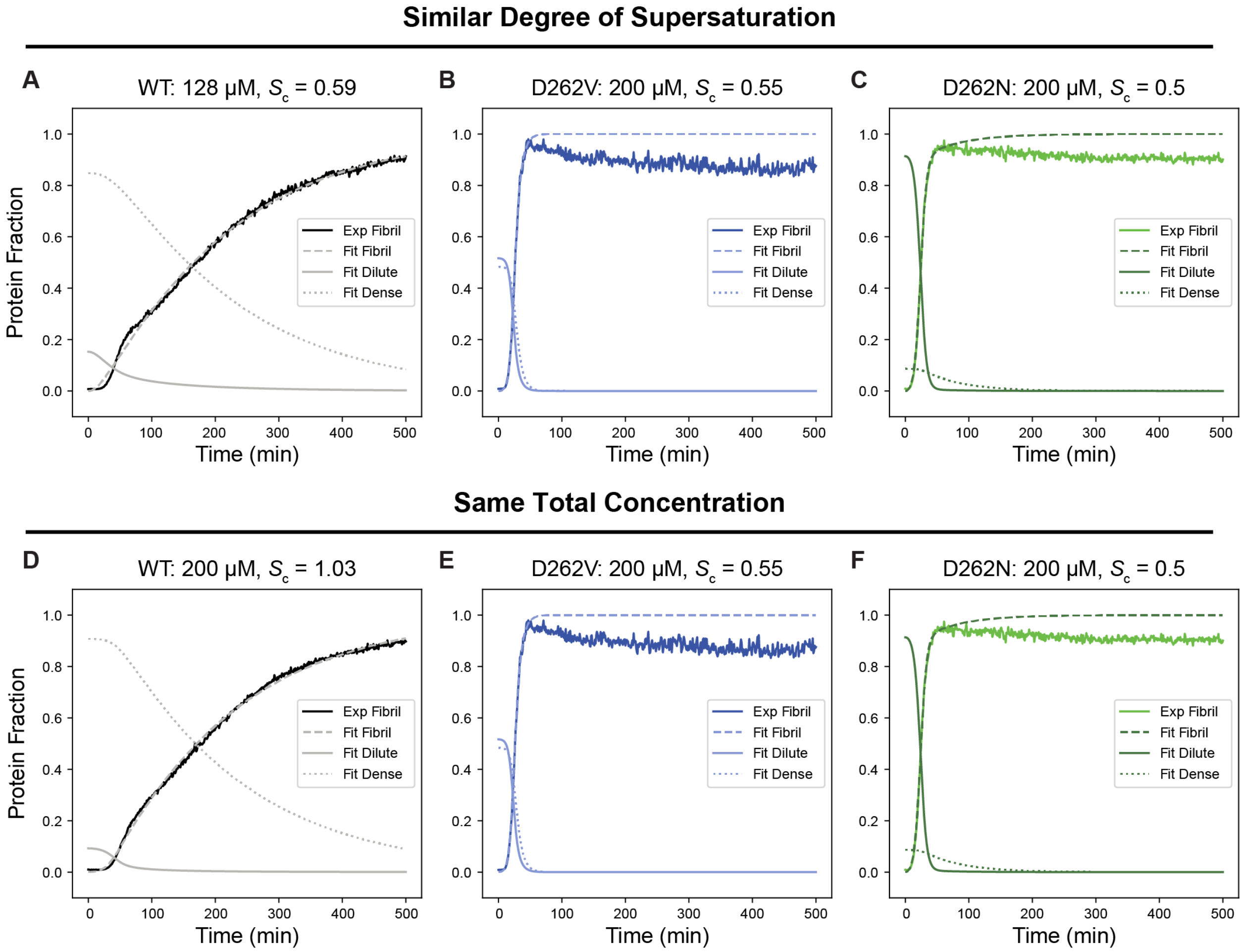


**Figure S5. Dense phase protein fraction of WT A1-LCD decays slowly leading to slower fibrillar growth, related to Figure 3.** The fraction of protein in the dilute, dense, and fibrillar phases as extracted using the chemical kinetics model for WT, D262V, and D262N at similar degrees of supersaturation (A-C) or same total concentration (D-F).

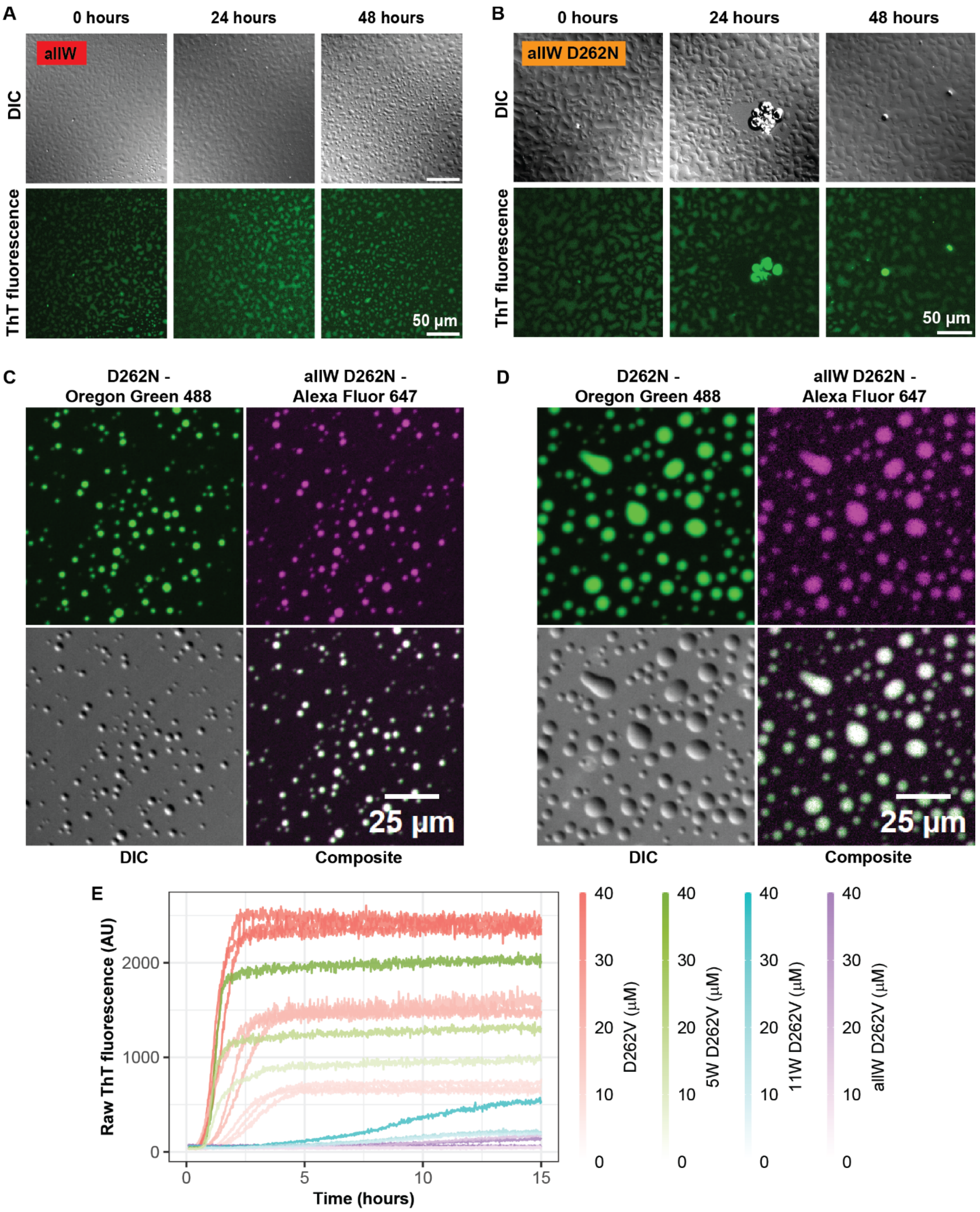


**Figure S6. allW variants form fibrils slowly and partition into A1-LCD condensates, related to Figure 4. (A, B)** DIC and fluorescence micrographs to monitor ThT fluorescence in samples with condensates formed by **(A)** allW and **(B)** allW D262N variants. **(C)** Colocalization of D262N (54 µM) and allW D262N (5 µM) in condensates. If each of the variants were alone in solution at these concentrations, D262N and allW D262N would be supersaturated. **(D)**

Colocalization of D262N (132 μM) and allW D262N (1 μM) in condensates. If each variant were the only variant in solution, allW D262N would be subsaturated at these concentrations and D262N would be supersaturated. (C, D) The labeling ratio of D262N with Oregon Green 488 and allW D262N with Alexa Fluor 647 were 1:100 and 1:300, respectively. **(E)** ThT fluorescence-monitored fibrillization kinetics of A1-LCD variants D262V, allW D262V, 11W D262V and 5W D262V at concentrations indicated by the color bar, showing raw ThT fluorescence as a function of time.

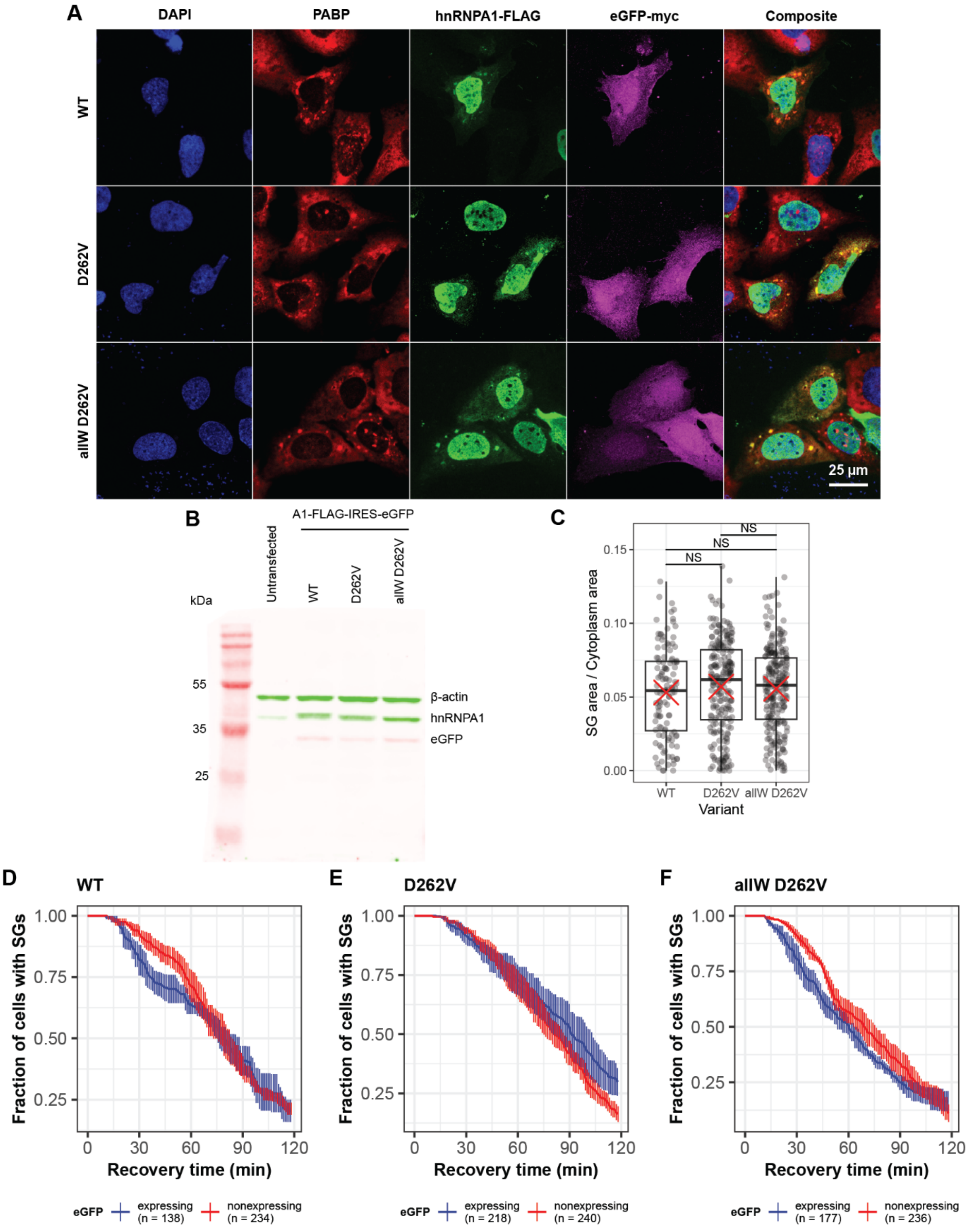


**Figure S7. hnRNPA1 constructs localize to stress granules, related to Figure 7. (A)** Immunofluorescence micrographs show colocalization of the stress granule marker PABP1 (red) and transiently transfected hnRNPA1-FLAG (green) in U2OS cells after 1 hour of heat stress at 43°C. The cell nuclei are stained with DAPI (blue). eGFP-expressing cells are shown as purple. Composite images combine only the red, blue and green channels. Scale bar, 25 µm. **(B)** Western blot showing expression of FLAG-tagged hnRNPA1 variants proportionately expressing eGFP via an IRES sequence. hnRNPA1 was detected using rabbit antibody raised against N-terminal region of hnRNPA1 (residues 8-42). Myc-tagged eGFP and β-actin were detected using respective antibodies raised in mouse and rabbit respectively. **(C)** Comparison between stress granule area fractions in eGFP-expressing cells at 17 minutes of heat stress. The box indicates the first and third quartiles and the central line indicates the second quartile. The whiskers indicate the range of the data. The means are shown as red crosses. NS: $p > 0.05$. **(D-F)** Comparison between SG disassembly kinetics between eGFP expressing and non-expressing cells for the indicated variants. The error bars indicate the SEM.

**Table S1. Sequences of WT A1-LCD and the variants used for in vitro experiments; related to Figures 1 and 4.**

| Variant | Amino acid sequence[a] |
|---|---|
| **WT** | *GS*MASASSSQRGRSGSGNFGGGRGGGFGGNDNFGRGGNFSGRGGFGGSRGGGGYG<br>GSGDGYNGFGNDGSNFGGGGSYNDFGNYNNQSSNFGPMKGGNFGGRSSGPYGGGG<br>QYFAKPRNQGGYGGSSSSSSYGSGRRF |
| **D262V** | *GS*MASASSSQRGRSGSGNFGGGRGGGFGGNDNFGRGGNFSGRGGFGGSRGGGGYG<br>GSGDGYNGFGNDGSNFGGGGSYNVFGNYNNQSSNFGPMKGGNFGGRSSGPYGGGG<br>QYFAKPRNQGGYGGSSSSSSYGSGRRF |
| **D262N** | *GS*MASASSSQRGRSGSGNFGGGRGGGFGGNDNFGRGGNFSGRGGFGGSRGGGGYG<br>GSGDGYNGFGNDGSNFGGGGSYNNFGNYNNQSSNFGPMKGGNFGGRSSGPYGGGG<br>QYFAKPRNQGGYGGSSSSSSYGSGRRF |
| **−4D+4V** | *GS*MASASSSQRGRSGSGNFGGGRGGGFGGNVNFGRGGNFSGRGGFGGSRGGGGYG<br>GSGVGYNGFGNVGSNFGGGGSYNVFGNYNNQSSNFGPMKGGNFGGRSSGPYGGGG<br>QYFAKPRNQGGYGGSSSSSSYGSGRRF |
| **−4D+4N** | *GS*MASASSSQRGRSGSGNFGGGRGGGFGGNNNFGRGGNFSGRGGFGGSRGGGGYG<br>GSGNGYNGFGNNGSNFGGGGSYNNFGNYNNQSSNFGPMKGGNFGGRSSGPYGGGG<br>QYFAKPRNQGGYGGSSSSSSYGSGRRF |
| **−3G1S+4V** | *GS*MASASSSQRGRSGSGNFGVGRGGGFGGNDNFGRGGNFSGRGGFGGVRGGGGYG<br>GSGDGYNGFGNDGSNFGGGGSYNDFGNYNNQSSNFVPMKGGNFGGRSSGPYGGGG<br>QYFAKPRNQVGYGGSSSSSSYGSGRRF |
| **−3D+3V** | *GS*MASASSSQRGRSGSGNFGGGRGGGFGGNVNFGRGGNFSGRGGFGGSRGGGGYG<br>GSGVGYNGFGNVGSNFGGGGSYNDFGNYNNQSSNFGPMKGGNFGGRSSGPYGGGG<br>QYFAKPRNQGGYGGSSSSSSYGSGRRF |
| **−3D+3N** | *GS*MASASSSQRGRSGSGNFGGGRGGGFGGNNNFGRGGNFSGRGGFGGSRGGGGYG<br>GSGNGYNGFGNNGSNFGGGGSYNDFGNYNNQSSNFGPMKGGNFGGRSSGPYGGGG<br>QYFAKPRNQGGYGGSSSSSSYGSGRRF |
| **allW** | *GS*MASASSSQRGRSGSGNWGGGRGGGWGGNDNWGRGGNWSGRGGWGGSRGGGGWG<br>GSGDGWNGWGNDGSNWGGGGSYNDFGNWNNQSSNWGPMKGGNWGGRSSGPWGGGG<br>QWWAKPRNQGGWGGSSSSSSWGSGRRW |
| **allW D262V** | *GS*MASASSSQRGRSGSGNWGGGRGGGWGGNDNWGRGGNWSGRGGWGGSRGGGGWG<br>GSGDGWNGWGNDGSNWGGGGSYNVFGNWNNQSSNWGPMKGGNWGGRSSGPWGGGG<br>QWWAKPRNQGGWGGSSSSSSWGSGRRW |
| **allW D262N** | *GS*MASASSSQRGRSGSGNWGGGRGGGWGGNDNWGRGGNWSGRGGWGGSRGGGGWG<br>GSGDGWNGWGNDGSNWGGGGSYNNFGNWNNQSSNWGPMKGGNWGGRSSGPWGGGG<br>QWWAKPRNQGGWGGSSSSSSWGSGRRW |
| **5WD262V** | *GS*MASASSSQRGRSGSGNFGGGRGGGFGGNDNFGRGGNWSGRGGWGGSRGGGGWG<br>GSGDGYNGFGNDGSNFGGGGSYNVFGNYNNQSSNFGPMKGGNFGGRSSGPYGGGG<br>QYFAKPRNQGGWGGSSSSSSYGSGRRW |
| **11WD262V** | *GS*MASASSSQRGRSGSGNWGGGRGGGWGGNDNWGRGGNWSGRGGWGGSRGGGGWG<br>GSGDGWNGWGNDGSNFGGGGSYNVFGNYNNQSSNFGPMKGGNFGGRSSGPYGGGG<br>QYFAKPRNQGGWGGSSSSSSWGSGRRW |

[a] Cleavage of the His-tag by TEV protease leaves an additional GS sequence at the N-terminus (shown in italics).

**Table S2. Amino acid sequences of full-length hnRNPA1 variants used in experiments in cells, related to Figure 7.**

| Variant | Amino acid sequence[a] |
|---|---|
| **WT** | MSKSESPKEPEQLRKLFIGGLSFETTDESLRSHFEQWGTLTDCVVMRDPNTKRSR<br>GFGFVTYATVEEVDAAMNARPHKVDGRVVEPKRAVSREDSQRPGAHLTVKKIFVG<br>GIKEDTEEHHLRDYFEQYGKIEVIEIMTDRGSGKKRGFAFVTFDDHDSVDKIVIQ<br>KYHTVNGHNCEVRKALSKQEMASASSSQRGRSGSGNFGGGRGGGFGGNDNFGRGG<br>NFSGRGGFGGSRGGGGYGGSGDGYNGFGNDGSNFGGGG**SYNDFG**NYNNQSSNFGP<br>MKGGNFGGRSSG**PY**GGGGQYFAKPRNQGGYGGSSSSSSYGSGRRF***DYKDDDDK*** |
| **D262V** | MSKSESPKEPEQLRKLFIGGLSFETTDESLRSHFEQWGTLTDCVVMRDPNTKRSR<br>GFGFVTYATVEEVDAAMNARPHKVDGRVVEPKRAVSREDSQRPGAHLTVKKIFVG<br>GIKEDTEEHHLRDYFEQYGKIEVIEIMTDRGSGKKRGFAFVTFDDHDSVDKIVIQ<br>KYHTVNGHNCEVRKALSKQEMASASSSQRGRSGSGNFGGGRGGGFGGNDNFGRGG<br>NFSGRGGFGGSRGGGGYGGSGDGYNGFGNDGSNFGGGG**SYNVFG**NYNNQSSNFGP<br>MKGGNFGGRSSG**PY**GGGGQYFAKPRNQGGYGGSSSSSSYGSGRRF***DYKDDDDK*** |
| **D262N** | MSKSESPKEPEQLRKLFIGGLSFETTDESLRSHFEQWGTLTDCVVMRDPNTKRSR<br>GFGFVTYATVEEVDAAMNARPHKVDGRVVEPKRAVSREDSQRPGAHLTVKKIFVG<br>GIKEDTEEHHLRDYFEQYGKIEVIEIMTDRGSGKKRGFAFVTFDDHDSVDKIVIQ<br>KYHTVNGHNCEVRKALSKQEMASASSSQRGRSGSGNFGGGRGGGFGGNDNFGRGG<br>NFSGRGGFGGSRGGGGYGGSGDGYNGFGNDGSNFGGGG**SYNNFG**NYNNQSSNFGP<br>MKGGNFGGRSSG**PY**GGGGQYFAKPRNQGGYGGSSSSSSYGSGRRF***DYKDDDDK*** |
| **allW** | MSKSESPKEPEQLRKLFIGGLSFETTDESLRSHFEQWGTLTDCVVMRDPNTKRSR<br>GFGFVTYATVEEVDAAMNARPHKVDGRVVEPKRAVSREDSQRPGAHLTVKKIFVG<br>GIKEDTEEHHLRDYFEQYGKIEVIEIMTDRGSGKKRGFAFVTFDDHDSVDKIVIQ<br>KYHTVNGHNCEVRKALSKQEMASASSSQRGRSGSGNWGGGRGGGWGGNDNWGRGG<br>NWSGRGGWGGSRGGGGWGGSGDGWNGWGNDGSNWGGGG**SYNDFG**NWNNQSSNWGP<br>MKGGNWGGRSSG**PY**GGGGQWWAKPRNQGGWGGSSSSSSWGSGRRW***DYKDDDDK*** |
| **allW D262V** | MSKSESPKEPEQLRKLFIGGLSFETTDESLRSHFEQWGTLTDCVVMRDPNTKRSR<br>GFGFVTYATVEEVDAAMNARPHKVDGRVVEPKRAVSREDSQRPGAHLTVKKIFVG<br>GIKEDTEEHHLRDYFEQYGKIEVIEIMTDRGSGKKRGFAFVTFDDHDSVDKIVIQ<br>KYHTVNGHNCEVRKALSKQEMASASSSQRGRSGSGNWGGGRGGGWGGNDNWGRGG<br>NWSGRGGWGGSRGGGGWGGSGDGWNGWGNDGSNWGGGG**SYNVFG**NWNNQSSNWGP<br>MKGGNWGGRSSG**PY**GGGGQWWAKPRNQGGWGGSSSSSSWGSGRRW***DYKDDDDK*** |
| **allW D262N** | MSKSESPKEPEQLRKLFIGGLSFETTDESLRSHFEQWGTLTDCVVMRDPNTKRSR<br>GFGFVTYATVEEVDAAMNARPHKVDGRVVEPKRAVSREDSQRPGAHLTVKKIFVG<br>GIKEDTEEHHLRDYFEQYGKIEVIEIMTDRGSGKKRGFAFVTFDDHDSVDKIVIQ<br>KYHTVNGHNCEVRKALSKQEMASASSSQRGRSGSGNWGGGRGGGWGGNDNWGRGG<br>NWSGRGGWGGSRGGGGWGGSGDGWNGWGNDGSNWGGGG**SYNNFG**NWNNQSSNWGP<br>MKGGNWGGRSSG**PY**GGGGQWWAKPRNQGGWGGSSSSSSWGSGRRW***DYKDDDDK*** |

[a] The C-terminal FLAG tag is shown in bold-italic. The LCD is underlined. The pathogenic mutation site (position 262) is denoted in red. The hexapeptide sequence near the pathogenic mutation site (positions 259-264) and the PY-NLS sequence (positions 288-289) are shown in bold. All mutation sites are highlighted in yellow.

#### Supplemental References


1. Bremer, A., Farag, M., Borcherds, W.M., Peran, I., Martin, E.W., Pappu, R.V., and Mittag, T. (2022). Deciphering how naturally occurring sequence features impact the phase behaviours of disordered prion-like domains. Nat Chem *14*, 196-207. 10.1038/s41557-021-00840-w.

2. Goldschmidt, L., Teng, P.K., Riek, R., and Eisenberg, D. (2010). Identifying the amylome, proteins capable of forming amyloid-like fibrils. Proceedings of the National Academy of Sciences *107*, 3487-3492. doi:10.1073/pnas.0915166107.
